# Genetic erosion reduces biomass temporal stability in wild fish populations

**DOI:** 10.1101/2019.12.20.884734

**Authors:** Jérôme G. Prunier, Mathieu Chevalier, Allan Raffard, Géraldine Loot, Nicolas Poulet, Simon Blanchet

## Abstract

Genetic diversity sustains species adaptation. However, it may also support key ecosystems functions and services that can be altered by the worldwide loss of genetic diversity. Here, using long-term wild fish data, we demonstrate that populations with high genetic diversity do not reach higher biomasses than populations with low genetic diversity. Nonetheless, populations with high genetic diversity have much more stable biomasses over recent decades than populations having suffered from genetic erosion, which has implications for the provision of ecosystem services and the risk of population extinction. Our results strengthen the importance of adopting prominent environmental policies to conserve this important biodiversity facet.

## INTRODUCTION

Biodiversity sustains critical ecosystem services, such as water filtering, pollination or biomass production^1^, that are directly compromised by the ongoing global biodiversity crisis^2^. By promoting trait complementarity or redundancy among species, interspecific diversity allows ecological communities to optimally capture essential resources, to transform those resources into biomass and to recycle them^3,4^. In species-rich communities, these ecological processes are maintained even in the face of environmental fluctuations, thus promoting ecosystem productivity and stability over time^5,6^: this is the insurance effect of species richness^7^.

Although biodiversity erosion is often associated to species loss, another form of erosion is silently underway: the loss of intraspecific genetic diversity^8^. Intraspecific genetic diversity can play a role similar to species diversity in driving ecological processes at the basis of ecosystem services, such as biomass production^9,10^. Beyond its positive influence on individual fitness and thus on *per capita* biomass production, intraspecific genetic diversity may favor functional complementarity or redundancy among individuals, thereby fostering a more efficient exploitation of available resources^1,7,9^. Genetically-diversified populations are therefore predicted to harbor both larger individuals (higher *per capita* biomass) and higher and more stable levels of total biomass than genetically-impoverished populations^9,10^. This direct relationship between intraspecific genetic diversity and biomass is expected to be particularly strong in ecosystems where species diversity is naturally low, which is actually the norm in many temperate ecosystems^11^. In such cases, the functioning of ecosystems probably depends more on the complementarity among genotypes than on the complementarity among species^12,13^, emphasizing the importance of maintaining genetic diversity to preserve ecosystem functions and services^9^.

Most studies investigating the relationship between intraspecific genetic diversity and key-ecological parameters such as biomass are based on experimental or semi-experimental settings, where population densities and levels of intraspecific genetic diversity are manipulated, while environmental conditions are controlled and maintained constant over time^9,14^. However, observational studies conducted in natural settings are still scarce and mostly concern plants^15^. Although these studies offer a number of advantages, experiments do not make it possible to cover large spatial and temporal scales or to investigate the influence of historical contingencies. Local levels of intraspecific genetic diversity indeed result from the interplay between long-term evolutionary trajectories (e.g., localization and size of glacial refugia^16^) and more recent –if not ongoing– ecological processes affecting individual life history traits or population demography (e.g., stressful environmental conditions^17^, bottleneck events^18^, or strong directional selection^19^). This natural complexity cannot be fully grasped by experimental studies. Observational field surveys are on the contrary more realistic and may provide important insights into the contribution of intraspecific genetic diversity, and the loss of it, to biomass and biomass stability in natural settings^15^. They yet raise several difficulties. First, assessing the influence of genetic diversity on biomass and biomass stability over several generations or seasonal cycles implies long-term monitoring programs of both population density and biomass, but such data are usually difficult to collect and are still scarce. Furthermore, the relationships between genetic diversity and biomass in across-population studies may be masked by the interplay with other factors also involved in biomass production, such as population density and environmental conditions, making it difficult to disentangle their respective contributions^15^. This last issue may however be partly alleviated through the use of causal modeling procedures (path analyses), making it possible to thoroughly confront theoretical expectations and experimental findings with the “real world”^3,20^.

Here, we capitalized on long-term field surveys of three parapatric non-commercial freshwater fish species (*Phoxinus dragarum, Gobio occitaniae* and *Squalius cephalus*) from two large river basins in south-western France (Fig. 1) to assess the relationships between intraspecific genetic diversity, total fish biomass and biomass temporal stability (measured as the inverse of biomass variability^21^), while controlling for the effect of environment (upstream-downstream gradient and eutrophication levels), of *per capita* biomass (or its temporal stability) and of past demographic events, using path analyses^22^. Total fish biomass stood for the total weight of all individuals from the three focal species collected at a given site (in g.m^-2^), standardized and then averaged across species and over years. Local levels of intraspecific genetic diversity were computed for each species using both microsatellite and SNPs data and similarly averaged across species. We addressed the following questions: Do the positive relationships found experimentally between intraspecific genetic diversity and total biomass and total biomass stability hold true in natural settings? If any, is the contribution of intraspecific genetic diversity to these ecosystem functions comparable in magnitude to that of other environmental determinants? Finally, is it possible to detect the impact of contemporary genetic erosion -i.e., the loss of intraspecific genetic diversity in response to a recent reduction in population size-on biomass and biomass stability of fish populations? This latter point is of high concern: with conservative estimates of 6-15% loss of intraspecific genetic diversity in wild organisms in the Anthropocene^23,24^, the impact of human-induced genetic erosion on natural ecosystems’ capacity to provide critical provisioning and regulating services to humanity may actually be much more important than anticipated.

**Figure 1.**
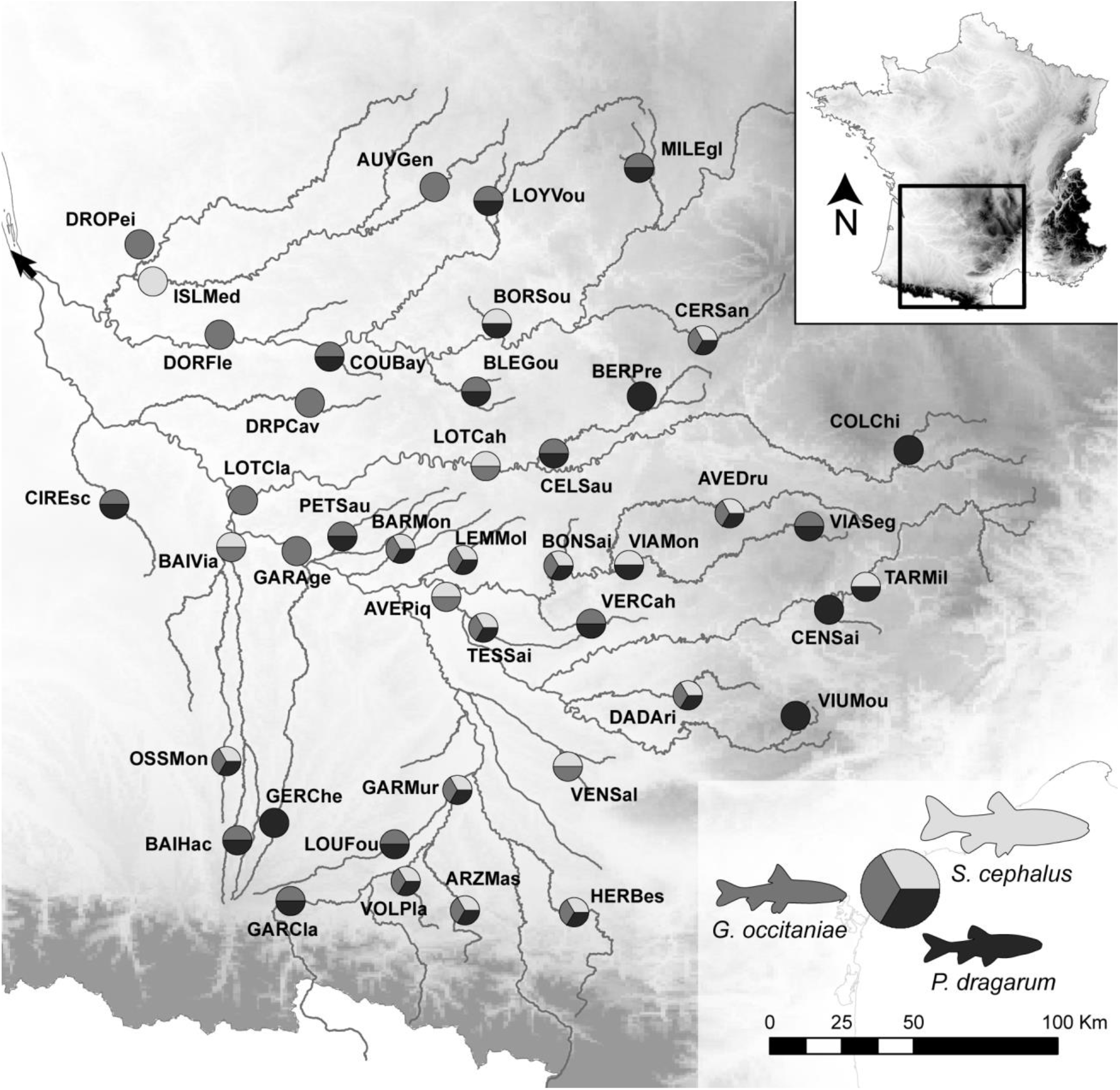
Study area and localization of sampled river stations and species. Geographic situation of the Garonne-Dordogne River basin in South-Western France and localization of the 42 unique river stations, with pie charts indicating species (co-)occurrence within each station. The black arrow indicates the location of the river mouth. Background is a shaded relief map.

## RESULTS

### Drivers of intraspecific genetic diversity

We found that intraspecific genetic diversity increased downstreamward (Fig. 2 and 3B; Supplementary Table 1), a classical pattern in rivers that could stem from asymmetrical gene flow, the presence of glacial refugees and/or higher effective population sizes in downstream areas^25^. Intraspecific genetic diversity was also indirectly impacted by water eutrophication, through a higher probability of having suffered from a bottleneck as eutrophication increases (Fig. 2 and 3A). As expected, the bottleneck probability altered spatial patterns of intraspecific genetic diversity: the loss of intraspecific genetic diversity associated with recent bottlenecks was particularly strong in downstream populations (Fig. 3B). This context-dependency of contemporary genetic erosion may reflect the observation that downstream areas are usually subject to multiple stressors (pollution, urbanization, channelization, …) that may reinforce the link between recent demographic changes and intraspecific genetic diversity.

**Figure 2.**
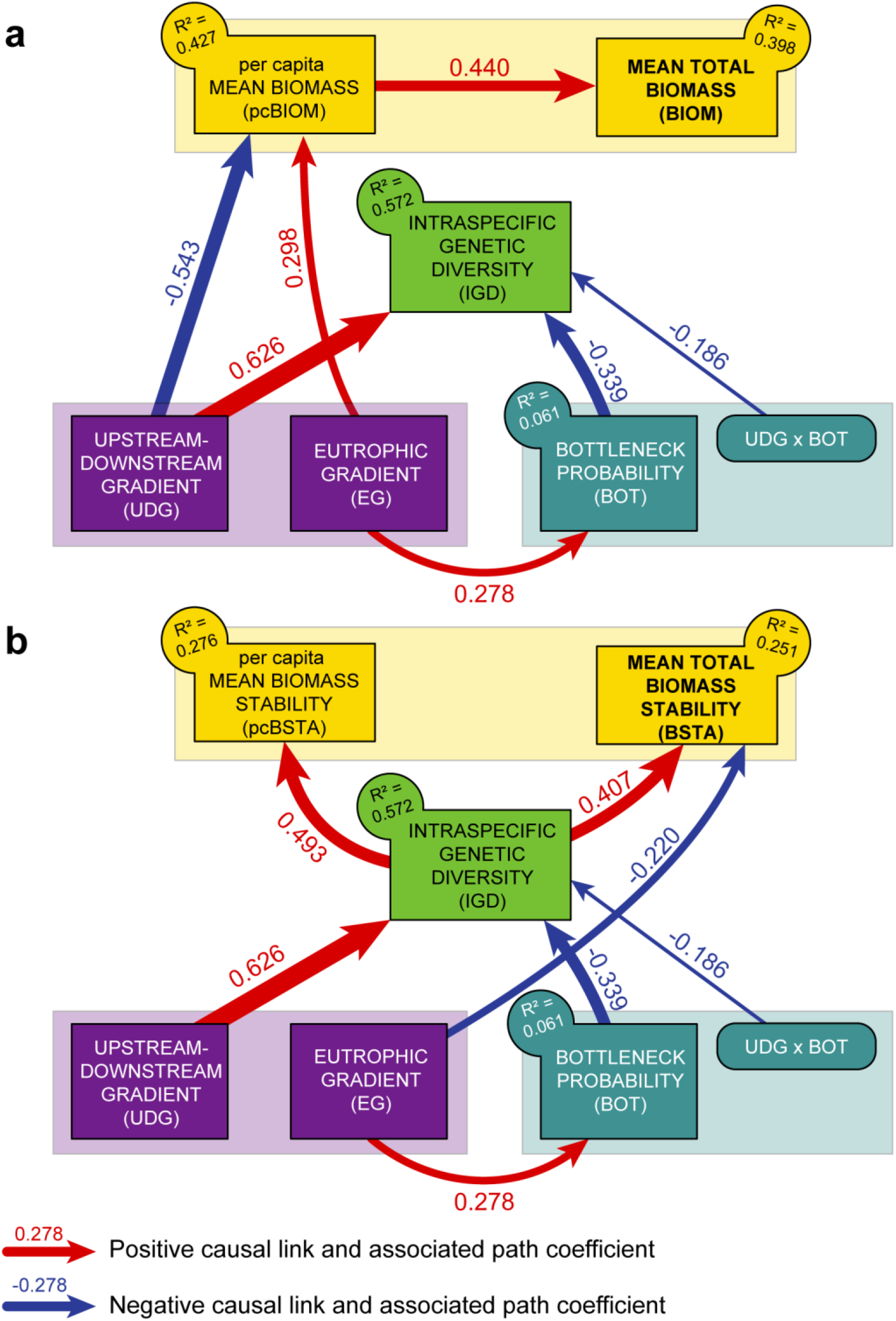
Final causal graphs. **a** Causal graph depicting the retained links among environmental (purple), bottleneck (blue), genetic (light green), and mean biomass variables (per capita and total; yellow). **b** Causal graph depicting the retained links among environmental (purple), bottleneck (blue), genetic (light green), and biomass stability variables (per capita and total; yellow). In each panel, retained first-order interactions are represented by rounded rectangles. Blue and red arrows represent negative and positive significant paths, respectively, with the width of arrows proportional to the absolute value of the corresponding path coefficient. Also provided is the amount of variance (R²) explained in each endogenous variable. Non-significant paths are not represented, for the sake of clarity (see Supplementary Table 1 for detailed results).

**Figure 3.**
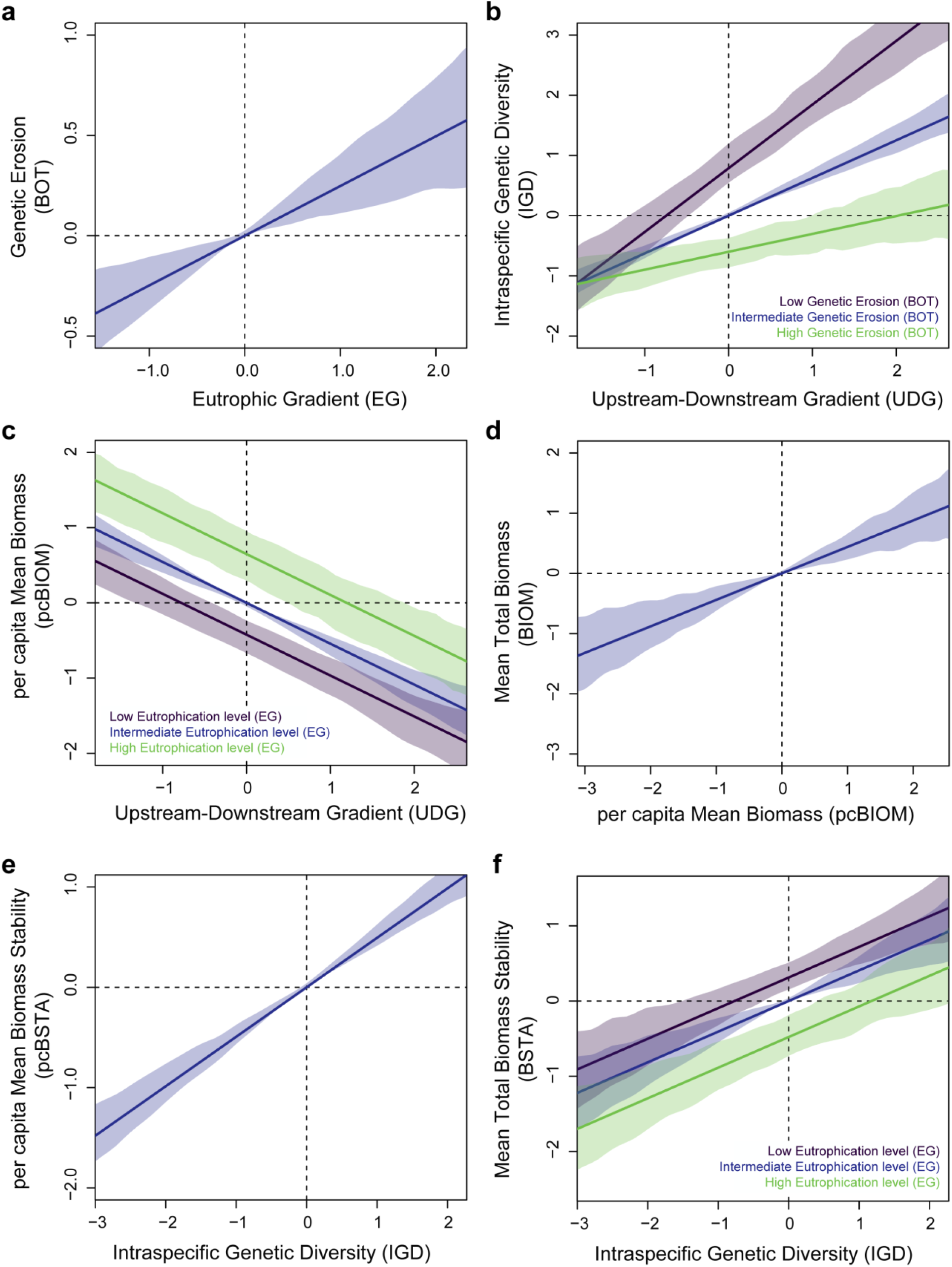
Predicted values and 95% confidence intervals of all endogenous variables given the retained links shown in Fig.1. When two predictors were considered (panels B, C and F), predictions were performed using the full range of the first predictor (x-axis) and the minimal (purple), mean (blue) and maximal (green) values of the second predictor.

### Drivers of total biomass

We found no significant relationship between intraspecific genetic diversity and either total biomass or *per capita* biomass (Fig. 2A; Supplementary Table 1). Yet, total biomass was directly linked to *per capita* biomass (Fig. 3D), indicating that total biomass stemmed from the presence of (a few) large individuals rather than that of many small individuals. *Per capita* biomass decreased downstreamward and increased with eutrophication (Fig. 3C), suggesting that, although eutrophication may indirectly have a long-term negative influence on populations and associated intraspecific genetic diversity, it may locally boost the overall system productivity by favoring individuals’ body condition^26^.

### Drivers of biomass stability

Contrastingly, we found that fish populations with higher levels of intraspecific genetic diversity displayed higher biomass stability over time than genetically-impoverished populations, whatever the temporal stability in *per capita* biomass (i.e., positive relationship between biomass stability and intraspecific genetic diversity: Fig. 2B and 3F; Supplementary Table 1). This important finding also held true when each species was analyzed separately (Supplementary Table 2). By favoring functional complementarity or redundancy among phenotypes^9,10,27^, higher genetic diversity likely allows populations to maintain an efficient exploitation of available resources in the face of natural environmental fluctuations, ensuring a stable production of biomass^1,7^. Biomass stability also tended to decrease with eutrophication (Fig. 2B and 3F; Supplementary Table 1), probably because biomass stability was negatively related to total biomass (Supplementary Note 1), and thus indirectly to *per capita* biomass, the latter increasing with eutrophication (Fig. 2A and 3). Fish biomasses were thus higher in the most eutrophic sites, but they were less stable over time: were this finding to be confirmed by further studies, this dual and opposite effect of eutrophication would have important implications for both the conservation of fish populations, since population stability is generally associated with lower extinction risk^28^. It is also noteworthy that temporal fluctuations in total fish biomass were unrelated to fluctuations in *per capita* biomass (that is, to temporal fluctuations in the average mass of individuals), reinforcing the hypothesis that biomass stability was rather indirectly fostered by mechanisms such as functional complementarity or redundancy among phenotypes^9,10,27^, acting as a biological insurance against natural environmental fluctuations^7^.

### Contribution of intraspecific genetic diversity to biomass stability

The contribution of intraspecific genetic diversity to the overall variance in biomass stability (R² = 21.1%) was much higher than that of considered environmental determinants (R² = 4%; Fig. 4). 84 % of the total explained variance in fish biomass stability was hence attributed to intraspecific genetic diversity, which suggests that intraspecific genetic diversity is a substantial driver of biomass stability in this area, although other unmeasured variables also likely sustain variation in biomass stability since a non-negligible part of this variation (∼75%; Fig. 4) remained unexplained by our model. Moreover, first-order interactions between intraspecific genetic diversity and environmental variables were not retained in final models (Fig. 2; but see the specific case of minnows in Supplementary Table 2), suggesting that the influence of intraspecific genetic diversity on total biomass stability may be predictable across environmental gradients, a result which is yet to be generalized to different taxa and ecosystems.

**Figure 4.**
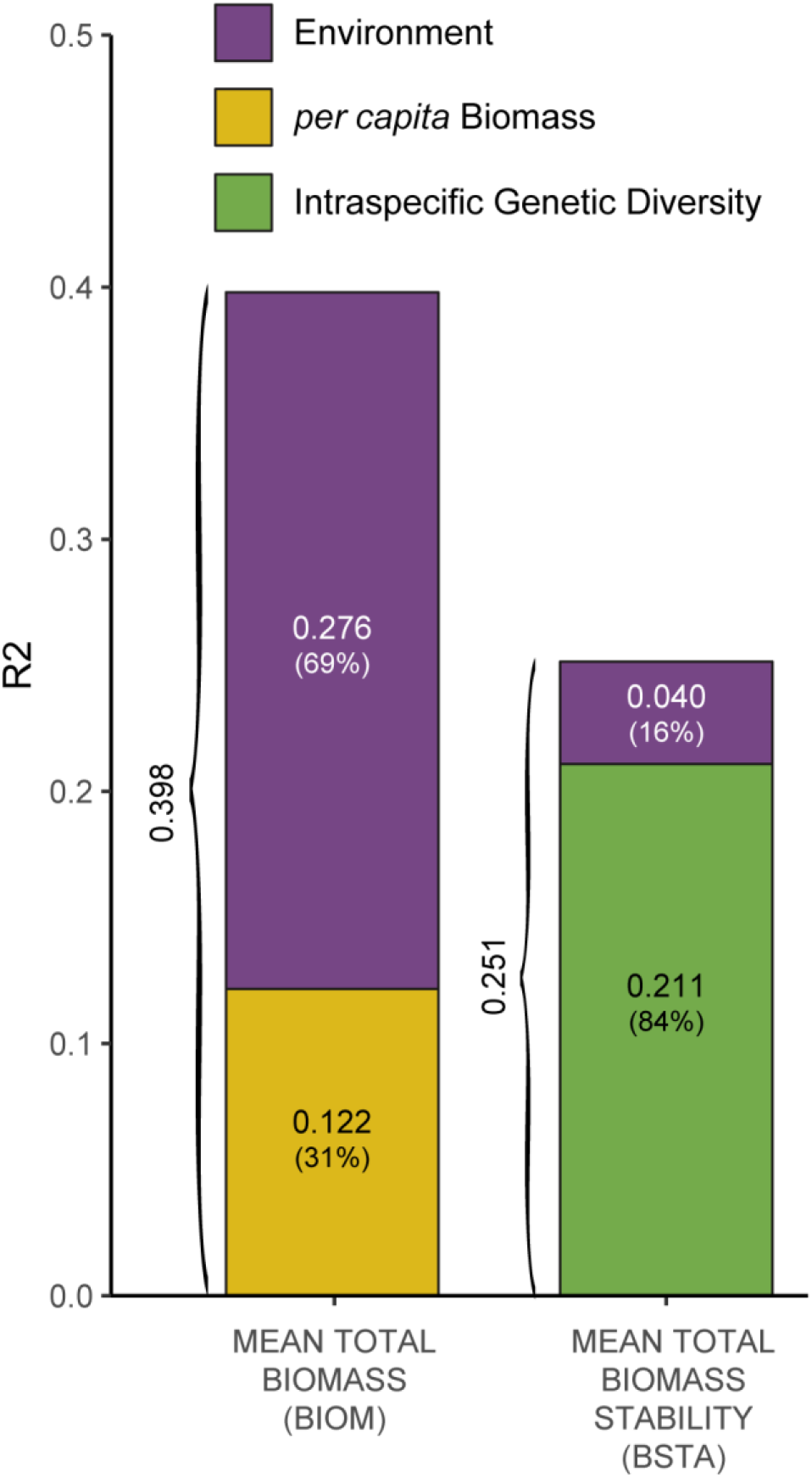
Variance partitioning. Contributions of environment variables (purple), *per capita* biomass variables (yellow) and intraspecific genetic diversity (light green) to the variance (R2) in Total Biomass (R² = 0.398) and Total Biomass stability (R² = 0.251). The contributions to the explained variance (in %) are indicated into brackets.

## DISCUSSION

Capitalizing on long-term demographic surveys, we report a positive relationship between intraspecific genetic diversity and temporal biomass stability in three freshwater fish species. This relationship indicates a buffering effect of intraspecific genetic diversity; genetically-impoverished populations being less efficient in maintaining stable biomass levels over time than genetically-diversified populations^7^. By favoring higher functional complementarity among phenotypes, higher genetic diversity likely allows populations to maintain an efficient exploitation of available resources in the face of natural environmental fluctuations, ensuring a stable production of biomass^1,4,7^. Interestingly, this buffering effect of intraspecific genetic diversity did not come with a performance-enhancing effect on biomass production^7^ (Fig. 2A): genetically-diversified populations did not show higher biomass levels compared to genetically-impoverished populations, mean total biomass being mostly driven by *per capita* biomass, and indirectly by the environment (Fig. 4). Nevertheless, our study provides one of the first non-experimental evidence that “real world” genetic diversity can directly promote temporal stability in biomass of wild organisms, in line with both theoretical expectations and experimental evidence^9,14^.

Our study being based on empirical data, it is not surprising that a large amount of variance in mean biomass and in biomass stability remained unexplained by our models (60 and 75% respectively; Fig. 2 and 4). Mean biomass and biomass stability were also probably driven by factors that we could not consider in this study, such as interspecific interactions at the community-level ^29^ or, at the ecosystem level, autotroph primary production ^30^ or terrestrial subsidies^31^. Nevertheless, intraspecific genetic diversity accounted for more than 21% of the variance in biomass stability, a contribution much higher than that of considered environmental predictors. Our findings not only indicate that the relationship between intraspecific genetic diversity and biomass stability holds true in natural ecosystems, but also that this relationship can be substantial compared to the effects of other undisputable determinants of biomass, as recently shown for interspecific diversity^5^. While species richness can buffer natural fish biomass production against environmental variations^20^, we argue that the intraspecific facet of biodiversity may actually also contribute to biomass stability in the wild ^9,14^.

A corollary to this finding is that the loss of intraspecific genetic diversity might undermine the temporal stability in biomass production. We notably detected a significant negative relationship between intraspecific genetic diversity and the probability that populations experienced a recent demographic contraction (i.e., recent bottleneck, Fig. 2), which is consistent with the imprint of recent human activities on contemporary levels of genetic diversity^23,24^. This finding must be considered with caution, since recent bottleneck probabilities were estimated using a modest number of microsatellite markers, which can generate bias in demographic inferences^32–35^. Specifically, the limited number of loci sampled in the genome^32^ and the departure from the assumed mutation model^33,34^ or from mutation-drift equilibrium^35^ generally increase the probability of inferring false signals of bottlenecks, i.e., of detecting a population decline in a truly stable population. This type of bias could have inflated the relationship we found between bottleneck probability and patterns of intraspecific genetic diversity. However, Paz-Vinas et al.^36^ demonstrated using simulations that, in river systems, demographic inferences based on microsatellite markers are rather robust to this bias, and actually more likely to detect false signals of expansion, i.e., detect a population expansion in a truly stable population, than to detect false signal of bottlenecks. Furthermore, to somehow limit the potential biases associated with the use of microsatellite markers, past bottleneck inferences were based on three independent methods that yielded congruent estimates (Supplementary Note 2). Although the links between past-demographic events and contemporary patterns of intraspecific genetic diversity would merit further confirmation based on genomic data^32^, we believe that it is reasonable to consider the low levels of intraspecific genetic diversity observed in some populations to stem -at least in part-from recent, possibly human-induced, demographic contractions, for instance triggered by water eutrophication (Fig. 2 and 3A). Although alternative and non-exclusive historical processes (e.g., post glacial colonization events) are also likely to explain observed spatial patterns of intraspecific genetic diversity^25,37^, this suggests that contemporary evolutionary processes can modulate ecological dynamics in natural settings. Since the loss of intraspecific genetic diversity always precede the loss of species^8^, genetic erosion may adversely affect key-ecological functions long before the first species of a community becomes extirpated. We therefore argue that the loss of intraspecific diversity observed worldwide^23,24^ may actually be responsible for a considerable alteration of many ecological processes in nature, but that these adverse effects might have been underestimated. With a loss of 6-15% in intraspecific genetic diversity^23,24^, we estimated a similar 8-10% reduction in biomass stability across the river basin (−8.9%; 95% confidence interval: [-10.1; -7.7]; Supplementary Note 3). This reduction in biomass stability was calculated using conservative estimates of intraspecific genetic diversity loss^23,24^; we therefore anticipate that this reduction could be much greater in species with weak conservation statuses, and that this reduction in stability may have important cascading effects on trophic networks, ecosystem functioning and, in some countries, food provisioning^1,38^.

Future studies are needed to confirm the significance of these results in other taxa and other ecosystems and to disentangle the relative contribution of intra- and interspecific diversity in explaining biomass production in the wild^10^. Nevertheless, our work suggests that the impact of genetic erosion on natural ecosystems’ capacity to provide critical provisioning and regulating services to humanity is probably more important than anticipated. This makes human-induced genetic erosion a critical conservation issue and stresses the need for human societies to adopt prominent environmental policies favoring all facets of biodiversity^39^.

## METHODS

### Sampling stations and biological models

We selected 47 river stations evenly scattered across two large River basins in South-Western France (the Garonne River basin and the Dordogne River basin) to reflect the environmental variability existing along the upstream–downstream gradients. Fish communities in these basins are generally poorly diverse (3 to 15 species^40^), and we focused on the three most common species: the minnow *Phoxinus dragarum*, the gudgeon *Gobio occitaniae* and the chub *Squalius cephalus*. These generalist cyprinids vary in their mean body length (minnows: 80–90 mm; gudgeons: 120–150 mm; chubs: 300–500 mm)^41^. They mainly feed on invertebrates (although chubs can also consume small-bodied fish) but occupy different habitats: chubs are primarily found in downstream sections at relatively low densities (∼0.01 ind.m^-2^), minnows are primarily found in upstream sections at relatively high densities (∼0.10 ind.m^-2^), whereas gudgeons are found all along the river basin in various habitats and at relatively high densities (∼0.08 ind.m^-2^)^41^. All stations are monitored yearly by the French Office for Biodiversity (OFB) with a constant sampling effort since 1990^42^. Demographic and biomass data from the three focal species were extracted from the OFB databases^43^. We only retained stations monitored from 1993 to 2020 with at least eight sampling sessions, resulting in 42 stations (Fig. 1; mean number of sampling sessions = 15.6; mean survey duration = 21.2 years; mean number of focal species per station = 2.0). The minimum pairwise distance among sampling stations was 18.5km (between TARMil and CENSai). This distance is higher than the maximum travelled distance recorded in chubs (16 km)^44^, here considered as the most mobile species, and all stations could thus be considered independent.

### Biomass data

For each species, station and year of survey, we collected local fish density (number of individuals per m²), total fish biomass (in g.m^-2^) and computed *per capita* biomass (or mean individual biomass, in g) as total fish biomass divided by local fish density. For each species, both total fish biomass and *per capita* biomass were standardized to make data comparable across species. For each station, we computed (i) Mean Total Biomass (respectively, *per capita* Mean Biomass) as the mean of total fish biomass (respectively, of *per capita* biomass) across species and over years, and to capture temporal fluctuations in biomass measures, (ii) Mean Total Biomass Stability (respectively, *per capita* Mean Biomass Stability) as the inverse of the squared coefficient of variation of Mean Total Biomass (respectively, of *per capita* Mean Biomass Stability) over years^45^. *Per capita* Biomass (stability) was here considered to determine whether Total Biomass (stability) was directly driven by the average individual biomass (stability) alone, or by other mechanisms such as functional complementarity among phenotypes. Each station was also assigned a sampling weight (ranging from 0.29 to 0.98) computed as the average of the relative local survey duration (compared to the maximal duration across stations) and of the relative local number of sampling sessions (compared to maximal number of sampling sessions across stations).

### Genetic data, intraspecific genetic diversity and bottleneck probability

The 42 retained stations were sampled in 2011 and 2014 with up to 30 adults from each species caught by electric-fishing. On the field, a small piece of pelvic fin was collected from each individual and was preserved in 70% ethanol, before releasing fish in situ. The genetic diversity of each population was characterized using both microsatellite (13 to 17 loci after filtering) and SNP markers (1244 to 1892 loci after filtering) as a composite metric extracted from both markers (see Supplementary Note 4 for details on DNA extraction, SNP calling, microsatellite genotyping and computation of metrics of genetic diversity). We used principal component analysis (PCA) to get a synthetic predictor of the cross-species level of intraspecific genetic diversity at each site. Only the first component was retained, accounting for 75.9 % of variance in genetic data, with genetically impoverished sites on the one hand (negative coordinates) and genetically diversified sites on the other hand (Supplementary Note 4). To determine the probability that populations suffered from a recent bottleneck, we used microsatellite data to compute three indirect yet complementary metrics of demographic contraction (using M-ratios^46^, VarEff^47^ and Migraine^48^), and used PCA to get a synthetic predictor of the cross-species local degree of genetic erosion (bottleneck probability) experienced by fish in recent generations. Only the first component was retained, accounting for 61.3 % of variance in input data, with sites with no sign of bottleneck on the one hand (negative coordinates) and sites having probably experienced a recent bottleneck on the other hand (Supplementary Note 2).

### Environment data

We similarly used PCA to get synthetic predictors of environment characteristics (distance to the river mouth, distance to the tributary source, water temperature and Water Quality Index) at the site level (Supplementary Note 5). The two first components were retained, accounting for 80.5 % of the total variance in environmental variables. The first component (55 % of variance) stood for the upstream-downstream gradient, with fresh upstream sites on the one hand (negative coordinates) and warmer downstream sites on the other hand. The second component (25.5 % of variance) stood for a eutrophic gradient, with oligotrophic river sites on the one hand (negative coordinates, low Water Quality Index) and nutrient-rich (mesotrophic) sites on the other hand (high Water quality Index).

### Path analyses

To investigate how intraspecific genetic diversity might influence Biomass and/or Biomass Stability while accounting for the effects of both environment and *per capita* Biomass^20,49^, we used path analyses^50,51^. We designed two full causal models describing the expected direct and indirect links among variables and their first-order interactions (Supplementary Note 6). Our main focus was on the direct links between intraspecific genetic diversity (intraspecific genetic diversity, or cross-product interactions of intraspecific genetic diversity with environmental variables^52^) and Total Biomass on the one hand (model A) and Biomass Stability on the other hand (model B). To consider the influence of *per capita* biomass on total biomass variables, we hypothesized that intraspecific genetic diversity would also indirectly promote Total Biomass (respectively, Biomass Stability), through a pathway involving *per capita* Biomass (respectively, *per capita* Biomass Stability). We further hypothesized that the environmental characteristics of stations (upstream-downstream gradient UDG, eutrophic gradient EG and the corresponding cross-product interaction UDGxEG) would affect *per capita* and total biomass variables both directly (for instance through higher intraspecific competition in harsh conditions) and indirectly, through pathways involving intraspecific genetic diversity (promoted for instance by higher proximity to glacial refugees^25^) as well as bottleneck (triggered for instance by pollutants). All variables were standardized to z-scores before using weighted path analyses^22^ with ‘MLM’ maximum likelihood estimation, ‘Satorra-Bentler’ scaled test statistic and station sampling weights. We simplified each model by removing non-significant paths one at a time, provided that cross-products were always associated with their additive terms^53^ and that removal led to an increase in the relative fit of the model (i.e., a decrease in BIC score^54^). The validity of final models was assessed according to their absolute fit (standardized root mean square residual (SRMR) < 0.09 and Robust Comparative Fit Index (CFI) > 0.96)^55^. Additionally, we verified that there was no significant discrepancy between the sample and the fitted covariance matrices^55^ (p-value associated with the model χ² statistic > 0.05). All path analyses were run using the R-package *lavaan*^56^. Predicted values of endogenous variables (Fig. 3) were obtained from linear regression models with each beta coefficient (and associated standard deviation) fixed to the value of the corresponding direct link within the final model. 95% confidence envelops were obtained by bootstrap (1000 iterations).

### Variance partitioning

For each total biomass variable, we computed (a) the amount of variance (R²) explained by each model. To assess the relative contribution of *per capita* biomass, intraspecific genetic diversity and environment to the variance in total biomass variables, we computing R² from further simplified models with (b) all variables related to *per capita* biomass being discarded (amount of variance explained by both environment and intraspecific genetic diversity), then (c) with all variables related to intraspecific genetic diversity (intraspecific genetic diversity and associated cross-products) being discarded (amount of variance explained by environment only). The relative contributions of *per capita* biomass and intraspecific genetic diversity to the variance in total biomass variables were respectively obtained by subtracting R² of (b) from R² of (a) and by subtracting R² of (c) from R² of (b).

## ACKNOWLEDGMENTS

This work was financially supported by the Office Français pour la Biodiversité (OFB) and the Agence Nationale pour la Recherche. We warmly thank all the colleagues and students who helped with field sampling and data analyses. The authors declare no competing interests.

## AUTHORS CONTRIBUTIONS

J.G.P., N.P. and S.B. formulated the main idea. J.G.P. collected environmental data and performed genetic/genomic analyses. M.C. and N.P. collected and prepared biomass data. J.G.P. performed data-analyses. J.G.P. and S.B. wrote the first draft and all authors provided feedback on subsequent versions of the manuscript.

## DATA ACCESSIBILITY

Raw data and data used to perform path analyses^57^: Figshare doi: 10.6084/m9.figshare.13095380.v8

## CODE ACCESSIBILITY

Code used to perform path analyses and produce figures^57^: Figshare doi: 10.6084/m9.figshare.13095380.v8

## REFERENCES

1. Cardinale, B. J. et al. Biodiversity loss and its impact on humanity. Nature 486, 59–67 (2012).

2. Barnosky, A. D. et al. Has the Earth’s sixth mass extinction already arrived? Nature 471, 51–57 (2011).

3. De Meester, L. et al. Analysing eco-evolutionary dynamics—The challenging complexity of the real world. Funct. Ecol. 33, 43–59 (2019).

4. Loreau, M. Biodiversity and ecosystem functioning: recent theoretical advances. Oikos 91, 3–17 (2000).

5. Duffy, J. E., Godwin, C. M. & Cardinale, B. J. Biodiversity effects in the wild are common and as strong as key drivers of productivity. Nature 549, 261–264 (2017).

6. Hooper, D. U. et al. Effects of Biodiversity on Ecosystem Functioning: A Consensus of Current Knowledge. Ecol. Monogr. 75, 3–35 (2005).

7. Yachi, S. & Loreau, M. Biodiversity and ecosystem productivity in a fluctuating environment: The insurance hypothesis. Proc. Natl. Acad. Sci. 96, 1463–1468 (1999).

8. Spielman, D., Brook, B. W. & Frankham, R. Most species are not driven to extinction before genetic factors impact them. Proc. Natl. Acad. Sci. 101, 15261–15264 (2004).

9. Hughes, A. R., Inouye, B. D., Johnson, M. T. J., Underwood, N. & Vellend, M. Ecological consequences of genetic diversity. Ecol. Lett. 11, 609–623 (2008).

10. Raffard, A., Santoul, F., Cucherousset, J. & Blanchet, S. The community and ecosystem consequences of intraspecific diversity: a meta-analysis: The ecological effects of intraspecific diversity. Biol. Rev. 94, 648–661 (2019).

11. Olson, D. M. et al. Terrestrial Ecoregions of the World: A New Map of Life on Earth. BioScience 51, 933 (2001).

12. Reusch, T. B. H., Ehlers, A., Hammerli, A. & Worm, B. Ecosystem recovery after climatic extremes enhanced by genotypic diversity. Proc. Natl. Acad. Sci. 102, 2826–2831 (2005).

13. Vellend, M. & Geber, M. A. Connections between species diversity and genetic diversity. Ecol. Lett. 8, 767–781 (2005).

14. Forsman, A. & Wennersten, L. Inter-individual variation promotes ecological success of populations and species: evidence from experimental and comparative studies. Ecography 39, 630–648 (2016).

15. Hendry, A. P. A critique for eco-evolutionary dynamics. Funct. Ecol. 33, 84–94 (2019).

16. Petit, R. J. Glacial Refugia: Hotspots But Not Melting Pots of Genetic Diversity. Science 300, 1563–1565 (2003).

17. Bestion, E. et al. Altered trophic interactions in warming climates: consequences for predator diet breadth and fitness. Proc. R. Soc. B Biol. Sci. 286, 20192227 (2019).

18. Jangjoo, M., Matter, S. F., Roland, J. & Keyghobadi, N. Connectivity rescues genetic diversity after a demographic bottleneck in a butterfly population network. Proc. Natl. Acad. Sci. 113, 10914–10919 (2016).

19. Coltman, D. W. et al. Undesirable evolutionary consequences of trophy hunting. Nature 426, 655–658 (2003).

20. Duffy, J. E., Lefcheck, J. S., Stuart-Smith, R. D., Navarrete, S. A. & Edgar, G. J. Biodiversity enhances reef fish biomass and resistance to climate change. Proc. Natl. Acad. Sci. 113, 6230–6235 (2016).

21. Donohue, I. et al. On the dimensionality of ecological stability. Ecol. Lett. 16, 421–429 (2013).

22. Grace, J. B. Structural equation modelling and natural systems. (Cambridge University Press, 2006).

23. Leigh, D. M., Hendry, A. P., Vázquez-Domínguez, E. & Friesen, V. L. Estimated six per cent loss of genetic variation in wild populations since the industrial revolution. Evol. Appl. 12, 1505–1512 (2019).

24. Exposito-Alonso, M. et al. Genetic diversity loss in the Anthropocene. Science 1431–1435 (2022).

25. Paz-Vinas, I., Loot, G., Stevens, V. M. & Blanchet, S. Evolutionary processes driving spatial patterns of intraspecific genetic diversity in river ecosystems. Mol. Ecol. 24, 4586–4604 (2015).

26. Hudon, C. et al. Oligotrophication from wetland epuration alters the riverine trophic network and carrying capacity for fish. Aquat. Sci. 74, 495–511 (2012).

27. Hughes, A. R. & Stachowicz, J. J. Genetic diversity enhances the resistance of a seagrass ecosystem to disturbance. Proc. Natl. Acad. Sci. 101, 8998–9002 (2004).

28. Finn, D. S., Bogan, M. T. & Lytle, D. A. Demographic Stability Metrics for Conservation Prioritization of Isolated Populations. Conserv. Biol. 23, 1185–1194 (2009).

29. Raffard, A., Santoul, F., Cucherousset, J. & Blanchet, S. The community and ecosystem consequences of intraspecific diversity: a meta-analysis: The ecological effects of intraspecific diversity. Biol. Rev. doi: 10.1111/brv.12472, (2019).

30. Delong, M. D. & Thorp, J. H. Significance of instream autotrophs in trophic dynamics of the Upper Mississippi River. Oecologia 147, 76–85 (2006).

31. Nakano, S. & Murakami, M. Reciprocal subsidies: Dynamic interdependence between terrestrial and aquatic food webs. Proc. Natl. Acad. Sci. 98, 166–170 (2001).

32. Hoban, S. M., Gaggiotti, O. E. & Bertorelle, G. The number of markers and samples needed for detecting bottlenecks under realistic scenarios, with and without recovery: a simulation-based study. Mol. Ecol. 22, 3444–3450 (2013).

33. Peery, M. Z. et al. Reliability of genetic bottleneck tests for detecting recent population declines. Mol. Ecol. 21, 3403–3418 (2012).

34. Putman, A. I. & Carbone, I. Challenges in analysis and interpretation of microsatellite data for population genetic studies. Ecol. Evol. 4399–4428 (2014) doi:10.1002/ece3.1305.

35. Chikhi, L., Sousa, V. C., Luisi, P., Goossens, B. & Beaumont, M. A. The confounding effects of population structure, genetic diversity and the sampling scheme on the detection and quantification of population size changes. Genetics 186, 983–995 (2010).

36. Paz-Vinas, I., Quéméré, E., Chikhi, L., Loot, G. & Blanchet, S. The demographic history of populations experiencing asymmetric gene flow: combining simulated and empirical data. Mol. Ecol. 22, 3279–3291 (2013).

37. De Kort, H. et al. Life history, climate and biogeography interactively affect worldwide genetic diversity of plant and animal populations. Nat. Commun. 12, 516 (2021).

38. Duffy, J. E., Lefcheck, J. S., Stuart-Smith, R. D., Navarrete, S. A. & Edgar, G. J. Biodiversity enhances reef fish biomass and resistance to climate change. Proc. Natl. Acad. Sci. 113, 6230–6235 (2016).

39. Laikre, L. et al. Post-2020 goals overlook genetic diversity. Science 367, 1083–1085 (2020).

40. Blanchet, S., Helmus, M. R., Brosse, S. & Grenouillet, G. Regional vs local drivers of phylogenetic and species diversity in stream fish communities. Freshw. Biol. 59, 450–462 (2014).

41. Keith, P., Persat, H., Feunteun, E., Adam, B. & Geniez, M. Les Poissons d’eau douce de France. (2011).

42. Poulet, N., Beaulaton, L. & Dembski, S. Time trends in fish populations in metropolitan France: insights from national monitoring data. J. Fish Biol. 79, 1436–1452 (2011).

43. Irz, P. et al. A long-term monitoring database on fish and crayfish species in French rivers. Knowl. Manag. Aquat. Ecosyst. 25 (2022) doi:10.1051/kmae/2022021.

44. Fredrich, F., Ohmann, S., Curio, B. & Kirschbaum, F. Spawning migrations of the chub in the River Spree, Germany. J. Fish Biol. 63, 710–723 (2003).

45. Haegeman, B. et al. Resilience, invariability, and ecological stability across levels of organization. http://biorxiv.org/lookup/doi/10.1101/085852 (2016) xdoi:10.1101/085852.

46. Garza, J. C. & Williamson, E. G. Detection of reduction in population size using data from microsatellite loci. Mol. Ecol. 10, 305–318 (2001).

47. Nikolic, N. & Chevalet, C. Detecting past changes of effective population size. Evol. Appl. 7, 663–681 (2014).

48. Leblois, R. et al. Maximum-Likelihood Inference of Population Size Contractions from Microsatellite Data. Mol. Biol. Evol. 31, 2805–2823 (2014).

49. Oehri, J., Schmid, B., Schaepman-Strub, G. & Niklaus, P. A. Biodiversity promotes primary productivity and growing season lengthening at the landscape scale. Proc. Natl. Acad. Sci. 114, 10160–10165 (2017).

50. Lefcheck, J. S. piecewiseSEM: Piecewise structural equation modeling in R for ecology, evolution, and systematics. Methods Ecol. Evol. 7, 573–579 (2015).

51. Fourtune, L. et al. Inferring Causalities in Landscape Genetics: An Extension of Wright’s Causal Modeling to Distance Matrices. Am. Nat. 191, 491–508 (2018).

52. Aiken, L. S., West, S. G. & Reno, R. R. Multiple regression: Testing and interpreting interactions. (SAGE Publications, 1991).

53. Batista-Foguet, J. M., Coenders, G., Saris, W. E. & Bisbe, J. Simultaneous Estimation of Indirect and Interaction Effects using Structural Equation Models. Metodološki Zv. 1, 163–184 (2004).

54. Burnham, K. P. & Anderson, D. R. Model selection and multimodel inference: a practical information-theoretic approach. (Springer, 2002).

55. Hu, L. & Bentler, P. M. Cutoff criteria for fit indexes in covariance structure analysis: Conventional criteria versus new alternatives. Struct. Equ. Model. Multidiscip. J. 6, 1–55 (1999).

56. Rosseel, Y. lavaan : An R Package for Structural Equation Modeling. J. Stat. Softw. 48, (2012).

57. Prunier, J. G. et al. Data for ‘Genetic erosion reduces biomass temporal stability in wild fish populations.’ (2022) doi:10.6084/m9.figshare.13095380.v8.

