## Supplementary material for "Genetic erosion reduces biomass temporal stability in wild fish populations"

**ORCIDs :** Jérôme G. Prunier 0000-0003-4110-2567, Simon Blanchet 0000-0002-3843-589X, Mathieu Chevalier 0000-0002-1170-5343

**Keywords**

Eco-evolutionary dynamics; Genetic erosion; Intraspecific diversity, Biodiversity-Ecosystem function.

**Supplementary Table 1.** Detailed results from simplified causal models.

For each model A (Mean Biomass) and B (Biomass Stability), the table provides absolute fit indices (CFI, SRMR, χ² statistic and associated two-tailed p-value; see Supplementary Note 7 for details), amounts of explained variance ($R^2$) in response variables and estimates of path coefficients with 95% confidence intervals obtained from bootstrap resampling (between square brackets) and associated statistics (z-values | p-values, between curly brackets). The significant positive relationship between IGD and Biomass stability is highlighted in bold. UDG: Upstream-downstream gradient; EG: Eutrophication gradient; IGD: Intraspecific genetic diversity; BOT: Bottleneck probability; BIOM: Total Biomass; BSTA: Total Biomass Stability; pcBIOM: *per capita* Mean Biomass; pcBSTA: *per capita* Mean Biomass Stability.

| Predictor | → Response | | **Model A** | **Model B** |
|---|---|---|---|---|
| | | | $\chi^2_{(14,42)}$ = 13.71, p-value = 0.471 | $\chi^2_{(14,42)}$ = 9.22, p-value = 0.817 |
| | | | CFI = 1 | CFI = 1 |
| | | | SRMR = 0.072 | SRMR = 0.075 |
| UDG | → BIOM | | -0.275 [-0.585; 0.034] / {-1.74 \| 0.081} | |
| pcBIOM | → BIOM | | 0.440 [0.083; 0.797] / {2.415 \| 0.016} | |
| | | $R^2$ | 0.398 | |
| **IGD** | **→ BSTA** | | | **0.407 [0.119; 0.695] / {2.772 \| 0.006}** |
| EG | → BSTA | | | -0.220 [-0.439; -0.001] / {-1.969 \| 0.049} |
| | | $R^2$ | | 0.251 |
| UDG | → pcBIOM | | -0.543 [-0.749; -0.338] / {-5.178 \| 0} | |
| EG | → pcBIOM | | 0.298 [0.032; 0.563] / {2.20 \| 0.028} | |
| | | $R^2$ | 0.427 | |
| IGD | → pcBSTA | | | 0.493 [0.325; 0.662] / {5.746 \| 0} |
| EG | → pcBSTA | | | -0.204 [-0.489; 0.081] / {-1.146 \| 0.161} |
| | | $R^2$ | | 0.276 |
| UDG | → IGD | | 0.626 [0.433; 0.820] / {6.341 \| 0} | |
| BOT | → IGD | | -0.339 [-0.576; -0.102] / {-2.804 \| 0.005} | |
| UDG x BOT | → IGD | | -0.186 [-0.346; -0.026] / {-2.276 \| 0.023} | |
| | | $R^2$ | 0.572 | |
| EG | → BOT | | 0.278 [0.006; 0.490] / {2.007 \| 0.045} | |
| | | $R^2$ | 0.061 | |

**Supplementary Table 2.** Detailed results from simplified causal models in chubs (A), gudgeons (B) and minnows (C and D). In each species, there is a positive relationship between IGD and BSTA, though mostly expressed in upstream areas in the case of minnows (D). See Supplementary Table 1 for legend.

(A) Results in chubs:

| Predictor | → Response | | Model A<br>$\chi^2_{(9,20)}$ = 4.78, p-value = 0.853<br>CFI = 1<br>SRMR = 0.086 | Model B<br>$\chi^2_{(11,20)}$ = 8.44, p-value = 0.750<br>CFI = 1<br>SRMR = 0.085 |
|---|---|---|---|---|
| pcBIOM | → BIOM | | 0.318 [-0.272; 0.909] / {1.057 \| 0.291} | |
| | | $R^2$ | 0.102 | |
| **IGD** | **→ BSTA** | | | **0.273 [0.020; 0.526] / {2.115 \| 0.034}** |
| pcBSTA | → BSTA | | | 0.645 [0.353; 0.938] / {4.321 \| 0} |
| EG | → BSTA | | | -0.242 [-0.562; 0.077] / {-1.487 \| 0.137} |
| | | $R^2$ | | 0.509 |
| UDG | → pcBIOM | | -0.474 [-0.815; -0.133] / {-2.727 \| 0.006} | |
| | | $R^2$ | 0.253 | |
| UDG | → pcBSTA | | | -0.242 [-0.503; 0.019] / {-1.816 \| 0.069} |
| | | $R^2$ | | 0.063 |
| UDG | → IGD | | 0.389 [0.105; 0.673] / {2.683 \| 0.007} | |
| BOT | → IGD | | -0.515 [-0.790; -0.240] / {-3.675 \| 0} | |
| | | $R^2$ | 0.684 | |
| UDG | → BOT | | -0.542 [-1.029; -0.055] / {-2.181 \| 0.029} | |
| | | $R^2$ | 0.284 | |

(B) Results in gudgeons:

| Predictor | → Response | | Model A<br>$\chi^2_{(13,33)}$ = 10.49, p-value = 0.653<br>CFI = 1<br>SRMR = 0.056 | Model B<br>$\chi^2_{(10,33)}$ = 5.26, p-value = 0.873<br>CFI = 1<br>SRMR = 0.063 |
|---|---|---|---|---|
| UDG | → BIOM | | -0.697 [-0.877; -0.517] / {-7.581 \| 0} | |
| EG | → BIOM | | 0.122 [-0.466; 0.333] / {1.13 \| 0.258} | |
| UDGxEG | → BIOM | | -0.289 [-0.272; -0.113] / {-3.215 \| 0.001} | |
| | | $R^2$ | 0.515 | |
| **IGD** | **→ BSTA** | | | **0.463 [0.186; 0.740] / {3.278 \| 0.001}** |
| pcBSTA | → BSTA | | | 0.265 [-0.009; 0.538] / {1.898 \| 0.058} |
| UDG | → BSTA | | | -0.581 [-0.856; -0.306] / {-4.143 \| 0} |
| | | $R^2$ | | 0.475 |
| UDG | → pcBIOM | | -0.612 [-0.782; -0.443] / {-7.084 \| 0} | |
| EG | → pcBIOM | | 0.240 [-0.023; 0.503] / {1.790 \| 0.073} | |
| | | $R^2$ | 0.512 | |
| UDG | → pcBSTA | | | -0.400 [-0.628; -0.172] / {-3.436 \| 0.001} |
| | | $R^2$ | | 0.200 |
| UDG | → IGD | | 0.506 [0.295; 0.717] / {4.702 \| 0} | |
| EG | → IGD | | 0.479 [0.272; 0.686] / {4.529 \| 0} | |
| BOT | → IGD | | -0.397 [-0.725; -0.069] / {-2.372 \| 0.018} | |
| | | $R^2$ | 0.563 | |
| EG | → BOT | | 0.166 [-0.182; 0.515] / {0.935 \| 0.35} | |
| | | $R^2$ | 0.028 | |

(C) Results in minnows:

| Predictor | → Response | | Model A | Model B |
|---|---|---|---|---|
| | | | **Model A** | **Model B** |
| | | | $\chi^2_{(13,31)}$ = 6.42, p-value = 0.930 | $\chi^2_{(15,31)}$ = 14.49, p-value = 0.489 |
| | | | CFI = 1 | CFI = 1 |
| | | | SRMR = 0.068 | SRMR = 0.073 |
| UDG | → BIOM | | -0.519 [-0.755; -0.284] / {-4.324 \| 0} | |
| pcBIOM | → BIOM | | 0.331 [0.157; 0.506] / {3.720 \| 0} | |
| | | $R^2$ | 0.460 | |
| pcBSTA | → BSTA | | | 0.256 [-0.084; 0.596] / {1.476 \| 0.14} |
| **UDG** | **→ BSTA** | | | **-0.445 [-0.929; 0.039] / {-1.803 \| 0.071}** |
| EG | → BSTA | | | -0.339 [-0.647; -0.030] / {-2.151 \| 0.031} |
| **IGD** | **→ BSTA** | | | **0.160 [-0.218; 0.538] / {0.831 \| 0.406}** |
| **UDGxIGD** | **→ BSTA** | | | **-0.367 [-0.685; -0.049] / {-2.265 \| 0.044}** |
| | | $R^2$ | | 0.305 |
| UDG | → pcBIOM | | -0.369 [-0.753; 0.015] / {-1.883 \| 0.06} | |
| EG | → pcBIOM | | 0.307 [-0.042; 0.656] / {1.725 \| 0.085} | |
| IGD | → pcBIOM | | 0.340 [-0.030; 0.710] / {1.801 \| 0.072} | |
| EGxIGD | → pcBIOM | | 0.438 [0.060; 0.815] / {2.270 \| 0.023} | |
| | | $R^2$ | 0.453 | |
| UDG | → pcBSTA | | | -0.028 [-0.337; 0.281] / {-0.178 \| 0.859} |
| EG | → pcBSTA | | | -0.010 [-0.431; 0.451] / {0.045 \| 0.964} |
| UDGxEG | → pcBSTA | | | 0.428 [0.053; 0.803] / {2.235 \| 0.025} |
| | | $R^2$ | | 0.109 |
| UDG | → IGD | | 0.421 [0.200; 0.641] / {3.740 \| 0} | |
| BOT | → IGD | | -0.230 [-0.535; 0.075] / {-1.477 \| 0.140} | |
| | | $R^2$ | 0.271 | |
| EG | → BOT | | 0.309 [-0.034; 0.652] / {1.767\| 0.077} | |
| | | $R^2$ | 0.094 | |

(D) Predicted values and 95% confidence intervals of BSTA in minnows given the retained links (pcBSTA and EG in panel A; first-order interaction between UDG and IGD in panel B) as indicated in Supplementary Table 2C. The positive relationship between IGD and BSTA is here mostly expressed in upstream areas.

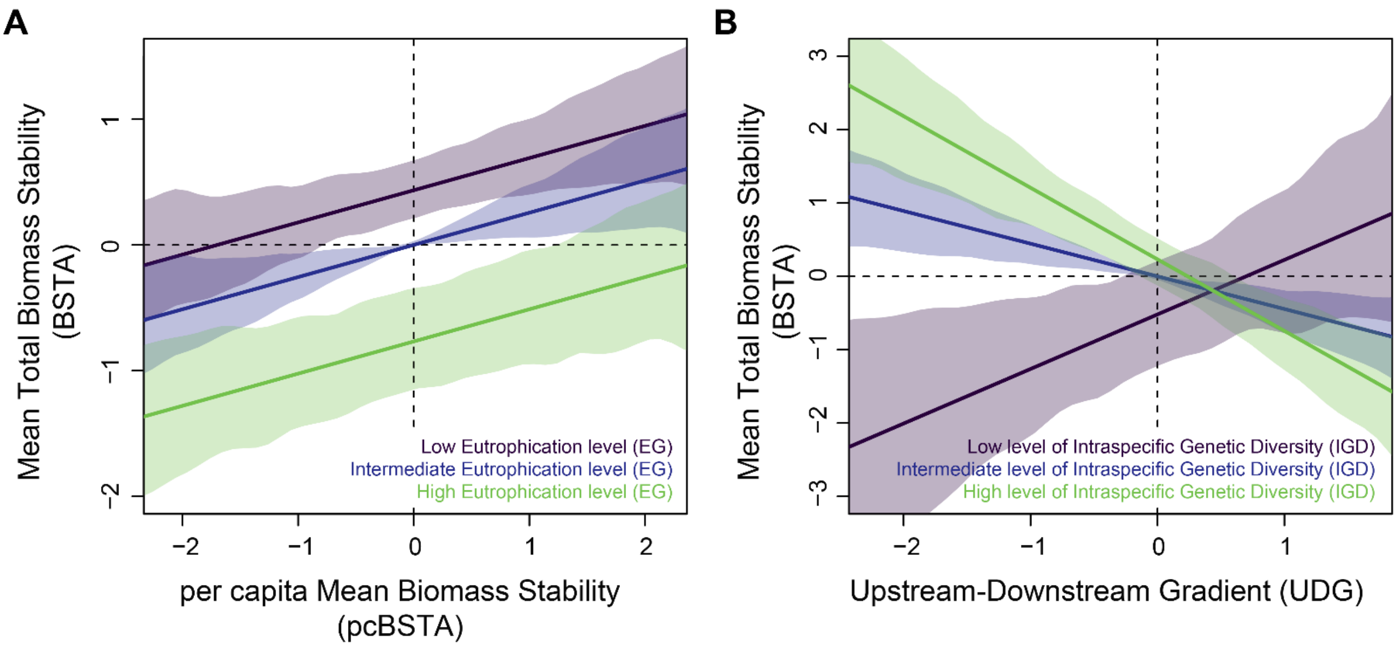

**Supplementary Note 1.** Demographic and biomass data.

Overall, Mean Total Biomass Stability (BSTA) was negatively related to Mean Total Biomass (BIOM), as illustrated in the figure below: the higher the Mean Total Biomass, the higher its fluctuations over the last decades.

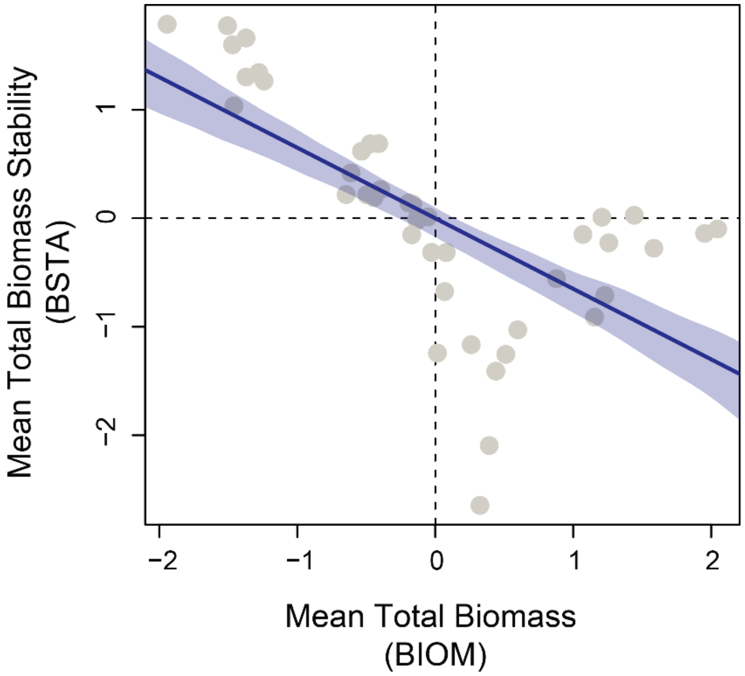

**Supplementary Note 2.** Bottleneck probability.

We used microsatellite data to compute three different quantitative measures of the degree of genetic erosion that populations underwent in recent generations: the M-ratio[24], the N-ratio computed with Migraine[25] (hereafter, MIratio) and the N-ratio computed with VarEff[26] (hereafter, VEratio). Note that the M-ratio can only detect signals of population decline (bottlenecks), but that both the MIratio and the VEratio can also detect signals of population expansion.

The M-ratio is the ratio between the number of observed alleles at a microsatellite locus and the allelic range of that locus, the latter being supposed to decrease slower than the number of alleles during a demographic collapse. This index, ranging from 0 to 1, is inversely proportional to the degree of genetic erosion[24], and has been shown to be particularly relevant in river systems[27]. For each station and each species, the M-ratio was computed for each microsatellite locus and then averaged over loci.

The MIratio was computed as $\theta_{cur}/\theta_{anc}$, that is the ratio between the scaled current population size $\theta_{cur}$ and the scaled ancestral population size $\theta_{anc}$ as inferred with Migraine[25]. The MIratio was estimated using the OnePopVarSize model, considering a single past change in population size, and a generalized stepwise mutation model (GSM). For each station and each species, PAC-likelihood computations were based on four iterations, 500 points and iteratively 2000 (first computation) or 20000 runs per point (second computation), to check consistency and improve convergence. In each case, we kept the estimate associated with the lowest RMS residual error, except when the algorithm failed to converge with the first computation, in which case we kept the second estimate.

The VEratio was similarly computed as $Ne_{cur}/Ne_{anc}$, that is the ratio between the estimated current effective population size $Ne_{cur}$ and the estimated ancestral effective population size $Ne_{anc}$ as inferred with the R-package VarEff[26]. For each station and each species, and following authors' recommendations, we first ran preliminary tests with short Markov chain Monte Carlo (MCMC) batches (1000 batches of length 1) to identify the best values for the number of past changes in effective population size (Jmax), for the effective size prior value (Nbar) and for the number of generations since the assumed origin of the population (Gbar). We then use these best values to run long MCMC batches (10000 batches of length 10), get effective population sizes at generation 1 ($Ne_{cur}$) and Gbar ($Ne_{anc}$), and compute VEratio. All runs were performed with 10 spaces between batches, a burnin period of 10000, a two-phase mutation model and a mutation rate of 0.0005.

MIratio and VEratio were log-transformed to meet normality assumptions. For each station, we averaged each metric over species and then used a principal component analysis (PCA) to get a synthetic predictor (bottleneck probability BOT) of the overall level of genetic erosion at the station level, with negative coordinates corresponding to stations with low genetic erosion (high M-ratio, MIratio and VEratio values). Only the first principal component (PC) was retained, accounting for 61.3 % of variance. The following table provides the minimal, maximal, mean and median values of raw metrics (out of 92 unique combinations of one species and one station), as well as their coordinates on the first axis and their contribution expressed in percentage. Note that a simulation study by Paz-Vinas et al.[27] showed that, in rivers, demographic inferences based on microsatellites are more likely to detect false signals of population expansion than false signals of population decline: here, we only detected 7 (7.6% of datasets) and 6 (6.5% of datasets) signals of expansion (log-ratio > 0) with the MIratio and the VEratio, respectively.

| | Min | Max | Mean | Median | Coordinates | Contribution |
|---|---|---|---|---|---|---|
| M-ratio | 0.427 | 0.880 | 0.697 | 0.705 | -0.871 | 41.23 |
| MIratio | -19.548 | 1.759 | -5.501 | -5.369 | -0.739 | 29.72 |
| VEratio | -4.489 | 1.031 | -1.694 | -1.518 | -0.731 | 29.05 |

The following figure provides a visual representation of eigenvalues (corresponding to the amount of the variation explained by each PC; panel A) and of the two first PC, with PC1 standing for bottleneck probability BOT.

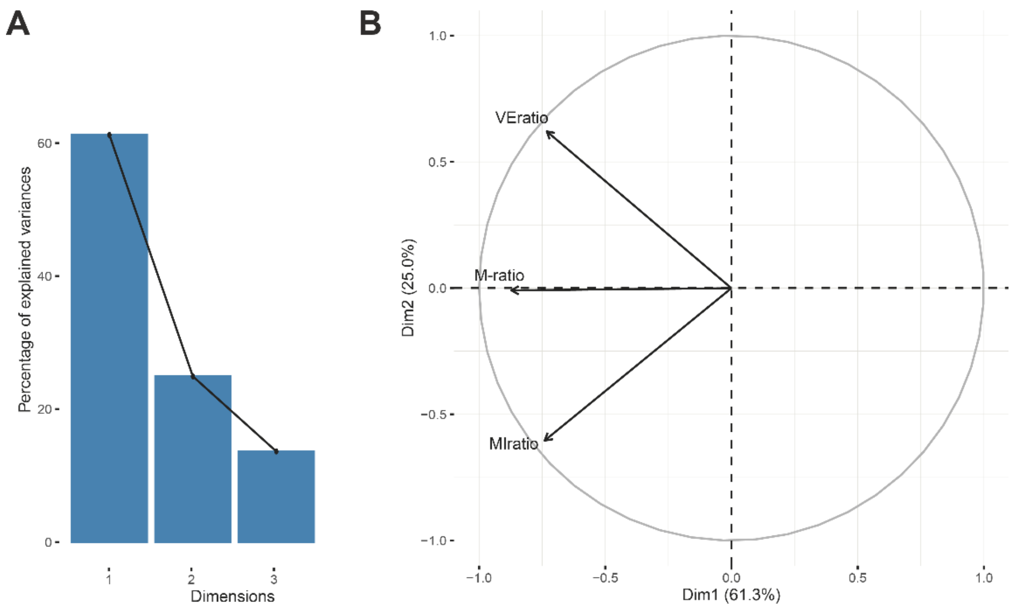

**Supplementary Note 3.** Overall expected change in biomass stability given a 6 to 15% decline in intraspecific genetic diversity.

We aimed to predict the overall expected change in biomass stability in our system given a 6 to 15% decline in intraspecific genetic diversity (IGD), as estimated by several authors in wild organisms[4,5]. Predictions were realized over k = 10000 iterations. For each iteration k, we first computed an eroded IGD predictor (eIGD) as:

$$eIGD_i = IGD_i - E_i \times \max(IGD)$$

with $IGD_i$ the observed IGD level at station i and $E_i$ an erosion factor randomly sampled from a uniform distribution ranging from 0.06 to 0.15.

For each iteration k, we then computed the predicted Biomass Stability BSTA from IGD and the predicted eroded Biomass Stability eBSTA from eIGD, using linear models and Eutrophication gradient EG as a covariate (considering final causal model B; see Fig. 1 in main text and Supplementary Table 1) as follows:

$$BSTA_{ki} = \beta_{ki}^{IGD\rightarrow BSTA} \times IGD_{ki} + \beta_{ki}^{EG\rightarrow BSTA} \times EG_{ki}$$

$$eBSTA_{ki} = \beta_{ki}^{IGD\rightarrow BSTA} \times eIGD_{ki} + \beta_{ki}^{EG\rightarrow BSTA} \times EG_{ki}$$

with $BSTA_{ki}$, $eBSTA_{ki}$, $EG_{ki}$, $IGD_{ki}$ and $eIGD_{ki}$ standing for the expected biomass stability, the expected eroded biomass stability, the observed eutrophic level, the observed IGD level and the previously computed eroded IGD level at station i, respectively; with $\beta_{ki}^{IGD\rightarrow BSTA}$ the effect of IGD on BSTA at station $\mathrm{i}$ as sampled from a normal distribution of mean μ=0.407 and σ=0.147 (corresponding to the 95% confidence interval of the path coefficient linking IGD to BSTA; Supplementary Table 1); with $\beta_{ki}^{EG\rightarrow BSTA}$ the effect of EG on BSTA at station $\mathrm{i}$ as sampled from a normal distribution of mean μ=-0.220 and σ=0.112 (corresponding to the 95% confidence interval of the path coefficient linking EG to BSTA; Supplementary Table 1).

For each iteration k, we finally collected the raw mean difference $D_k$ between the predicted eroded Biomass Stability eBSTA and the predicted Biomass Stability BSTA as:

$$D_k = \overline{eBSTA_{ki}} - \overline{BSTA_{ki}}$$

The predicted overall expected change in biomass stability given a 6 to 15% decline in IGD was finally computed as $\overline{D_k}$, with 95% quantiles as confidence interval.

The following histogram provides the distribution of $D_k$ values along with the corresponding fitted normal curve (in orange), mean value and 95% confidence intervals. We found $\overline{D_k}$ = -8.87 % [-10.11; -7.72].

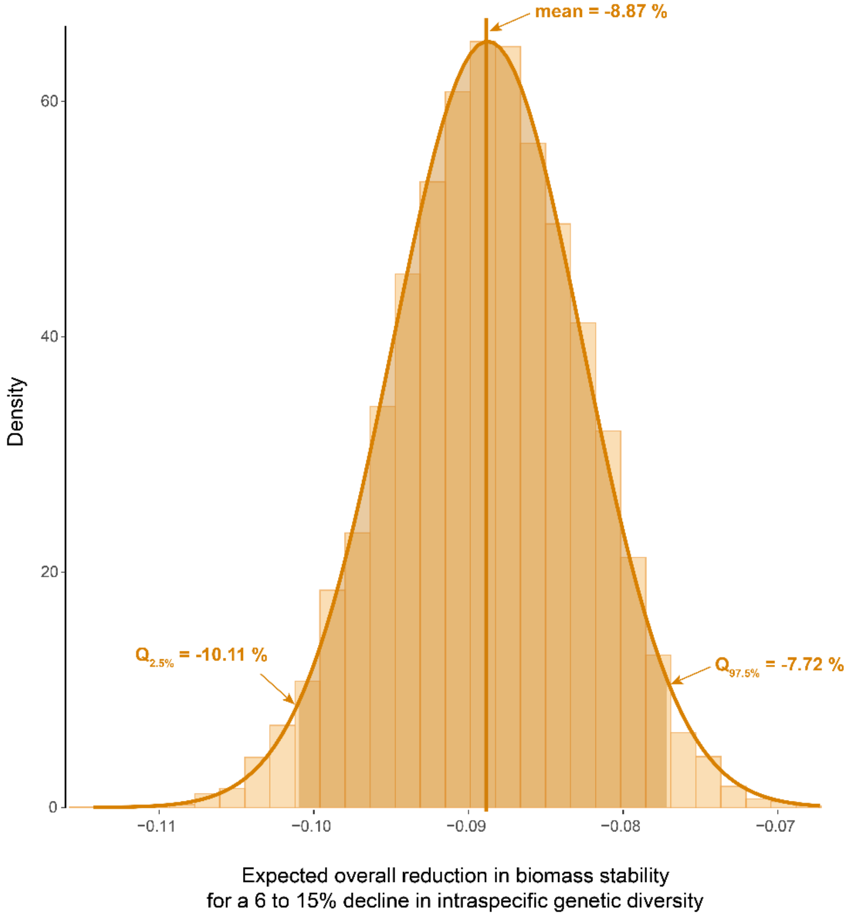

#### **Supplementary Note 4.** Genetic data.

**Field sampling**

The 42 stations were sampled in 2011 and in 2014 with up to 30 adults from each species caught by electric-fishing, resulting in a set of 35, 37 and 21 sampled populations in *P. dragarum*, *G. occitaniae* and *S. cephalus*, respectively. On the field, a small piece of pelvic fin was collected from each individual and was preserved in 70% ethanol, before releasing fish in situ. Genomic DNA was extracted using a salt-extraction protocol[6] and used to obtain, for each species, population-based SNP allelic frequencies following a paired-end pool-seq procedure[7] from material collected in 2014 (see "SNP data") as well as individual-based microsatellite genotypes from material collected in 2011 (see "Microsatellite data").

**SNP data**

For each species and each station, DNA from all individuals was pooled at equimolar concentrations to reach a total amount of 5mg of DNA. Individual concentrations were determined using a QuBit 2.0 fluorometer (2.0, Life Technologies, Carlsbad, CA, USA). In *P. dragarum* and *G. occitaniae*, pooled DNA from each population was homogenized and split into two replicates, for subsequent analysis of allelic frequencies reliability. Pooled DNA was digested as in [8] using SbfI restriction enzymes, followed by barcode ligation, sample pooling, DNA shearing, size selection of RAD tags (150 bp), adaptor ligation, RAD tag amplification and sequencing on two Hiseq lanes (GeT Platform, Toulouse, France). The procedure resulted in demultiplexed paired-end short reads that were subsequently processed for SNP identification and allelic frequencies estimation. All paired raw *fastq* files were filtered using the *process_radtags* and the *clone_filter* functions from Stacks[9], in order to remove reads with uncalled bases or low quality scores and discard PCR duplicates.

For each species, a single *fastq* file, corresponding to an outlier population as identified from microsatellites data (see "Outlier populations"), was then processed using the Velvet *de novo* sequence assembler [10] to design a draft reference genome. Velvet's assembly parameters were optimized using the VelvetOptimiser wrapper (http://bioinformatics.net.au/software.shtml), with 19 and 99 as starting and ending hash values, a minimum contig length of 150 pb and an insert length of 240pb. Draft genomes (in *fasta* format) were then indexed using both the *index* function from bwa [11] and the *faidx* function from SamTools [12].

All filtered paired-end *fastq* files were aligned on their draft genome using the *aln* and *sampe* functions from bwa. Aligned SAM files were converted to BAM format with the *view* and *sort* functions from SamTools, and filtered for unpaired, unmapped or badly mapped reads (mapping quality score < 20) using the *filter* function from BamTools [13]. For each species, all indexed and filtered BAM files were then assembled in a single *mpileup* file using the *mpileup* function from SamTools. These *mpileup* files were synchronized in Popoolation2[14] with the *mpileup2sync.jar* java script. SNP allelic frequencies were finally determined using the *snp-frequency-diff.pl* perl script in Popoolation2 with a minimum allele count of 4 and a coverage ranging from 30 to 400.

The whole procedure led to the identification of 10137, 13671 and 5897 SNPs in *P. dragarum*, *G. occitaniae* and *S. cephalus*, respectively. In *P. dragarum* and *G. occitaniae*, allelic frequencies reliability was assessed for each SNP and each station by comparing allelic frequencies between pairwise replicates. When allelic frequencies were available for the two replicates and when $\Delta_{AF}$, the difference in allelic frequencies between pairwise replicates, was lower than 0.25, the final allelic frequency was computed as the average of allelic frequencies across replicates. Otherwise, the final allelic frequency was set as missing data. In each species, we finally followed a two-step filtering procedure: (i) we first discarded any SNP with available allelic frequencies for less than 15 stations; (ii) we then discarded any station with available allelic frequencies for less than 150 SNPs. This final filtering procedure generated a total of 1244, 1892 and 1847 SNPs in *P. dragarum*, *G. occitaniae* and *S. cephalus*, respectively. The characteristics of final datasets are provided in table below.

| | *P. dragarum* | *G. occitaniae* | *S. cephalus* |
|---|---|---|---|
| Number of SNPs | 1244 | 1892 | 1847 |
| Number of populations | 31 | 34 | 17 |
| Mean coverage (± standard deviation) | 99.2 ± 41.9 | 77.5 ± 34.8 | 51 ± 13.1 |
| Percentage of missing AF values (across SNPs and populations) | 1.2 | 13.2 | 19.1 |

**Microsatellite data**

Genetic material collected in 2011 was used to genotype individuals at 18, 15 and 19 microsatellites markers in *P. dragarum*, *G. occitaniae* and *S. cephalus*, respectively. Polymerase chain reactions (PCR) and genotyping were performed as detailed in the supplementary file "Microsatellite PCR conditions.xlsx", resulting in a final dataset of 3262 genotypes (1177 in *P. dragarum*, 1227 in *G. occitaniae* and 858 in *S. cephalus*). We checked for multi-locus deviation from Hardy-Weinberg Equilibrium (HWE) and for gametic disequilibrium using GENEPOP 4.2.1[15] after sequential Bonferroni correction to account for multiple related tests[16]. In each species, the presence of null alleles was assessed by analyzing homozygote excess at each locus in five populations previously identified as panmictic, using MICROCHECKER 2.2.3[17]. We discarded from further analyses any locus showing significant gametic disequilibrium and/or evidence of null alleles, resulting in the withdrawal of one locus (CtoG-075) in *P. dragarum*, two loci (Lsou5 and Gob12) in *G. occitaniae* and three loci (Ca1, Lid11 and LleC-090) in *S. cephalus*, for a total number of 17, 13 and 16 loci in each species, respectively. Although the three focal species are of limited interest for anglers[18], discriminant analyses of principal components (dAPC[19]) performed on microsatellite data allowed identifying outlier populations, possibly resulting from past stocking events[20]. All outlier populations (one in *P. dragarum*, four in *G. occitaniae* and one in *S. cephalus*; see "Outlier populations") were subsequently discarded from further analyses. For each species, a single outlier population was yet considered for *de novo* genome assembly (see “SNP data”).

**Outlier populations**

A discriminant analysis of principal components (dAPC) was performed on each microsatellite dataset using the R-package *adegenet*[21]. In the following figure, arrows indicate outliers populations, probably resulting from past stocking events[20]. These populations were discarded from the final datasets. Asterisks indicate outlier populations used to design a draft reference genome in each species (see “SNP data”).

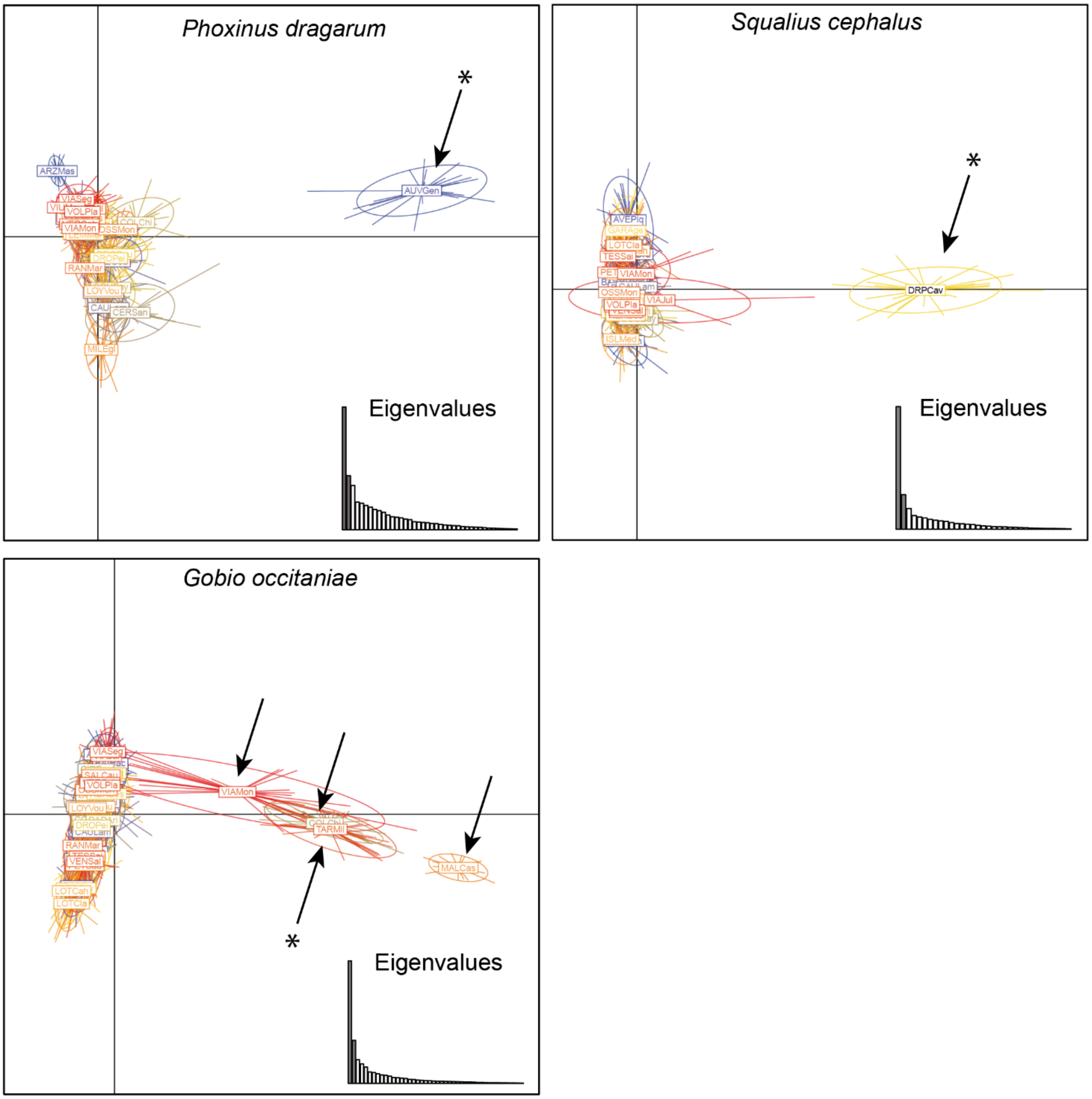

**Metrics of genetic diversity and characteristics of the first principal component based on genetic data.**

For each species and station, we computed a total of four metrics of genetic diversity. First, we used SNPs allelic frequencies to compute two metrics in R[22]: the expected level of heterozygosity across SNPs loci (sHe) and the observed level of SNP polymorphism (sPo), computed as the number of non-fixed loci (0 < allelic frequency < 1) divided by the total number of loci with non-missing data in a given population. We then used microsatellite data to compute two additional metrics, the expected (µHe) and observed (µHo) levels of heterozygosity across microsatellite loci, using the software GENETIX 4.3 [23].

These four metrics of genetic diversity naturally range between 0 and 1 and are thus directly comparable across species: for each station, we thus averaged each metric over species and then used a principal component analysis (PCA) to get a synthetic predictor of the overall level of genetic diversity at the station level. Only the first principal component (PC) was retained, accounting for 75.9 % of variance in genetic data. The following table provides the coordinates of raw variables on the first axis, as well their contribution expressed in percentage. µ: microsatellites / s: SNPs; He: expected heterozygosity / Ho: observed heterozygosity / Po: Polymorphism (see main text for details).

| | Coordinates | Contribution |
|---|---|---|
| µHe | 0.894 | 26.32 |
| µHo | 0.872 | 25.02 |
| sHe | 0.822 | 22.26 |
| sPo | 0.895 | 26.40 |

The following figure provides a visual representation of eigenvalues (corresponding to the amount of the variation explained by each PC; panel A) and of the two first PC, with PC1 standing for Intraspecific Genetic Diversity IGD. The percentage of variance explained by each component is also indicated.

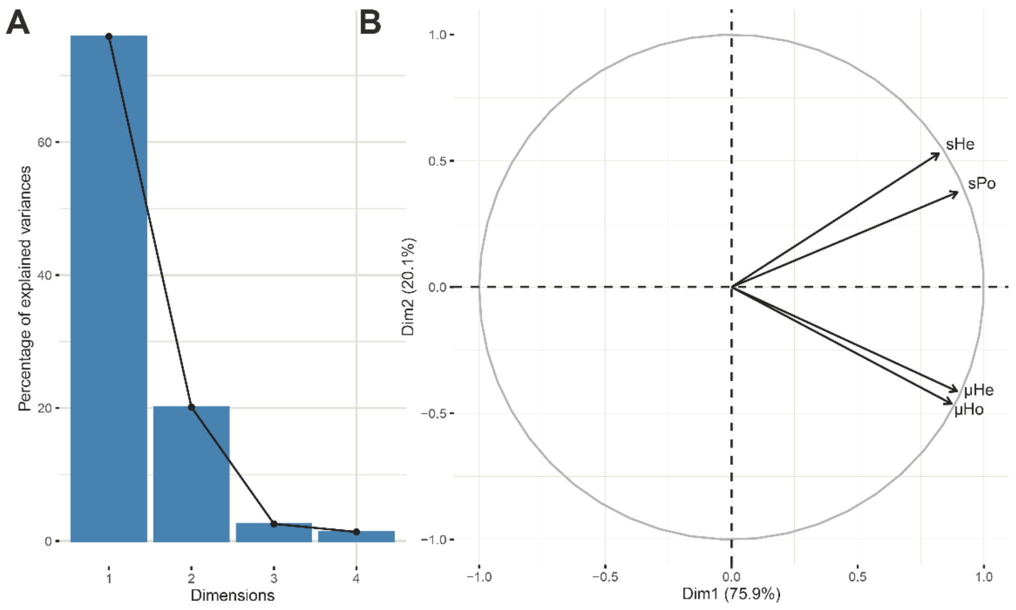
A
B
Percentage of explained variances
Dimensions
Dim2 (20.1%)
Dim1 (75.9%)
sHe
sPo
μHe
μHo


248

#### **Supplementary Note 5.** Environmental data.

##### Data collection and Water Quality Index

From the French Theoretical Hydrological Network[28], we computed the distance to the mouth (in m) and the distance to the source (in m). From the database of the Water Information System of the Adour Garonne basin (http://adour-garonne.eaufrance.fr), we collected ten additional variables related to water quality, measured in June, July and August from 2000 to 2015: Temperature (in °C), oxygen concentration (in $mg.L^{-1}$) and saturation (in %), Biochemical oxygen demand (in $mg.L^{-1}$), as well as concentrations (in $mg.L^{-1}$) in nitrogen compounds (ammonium $NH^{4}+$, nitrates $NO^{3-}$ and nitrites $NO^{2-}$), in phosphorus compounds (total phosphorus P and phosphate $PO_4^{3-}$) and in dissolved organic carbon. Following criteria used by French managers to assess the ecological status of rivers from various physico-chemical parameters according to the French implementation of the European Water Framework Directive 2000/60/EC (see “Distribution of main chemical components concentrations in the 47 river stations”), we assigned to each station, each month of survey and each water quality variable but temperature a value ranging from 1 (very good water quality) to 6 (very bad water quality). For each station, values were then averaged over water quality variables, then over months and finally over years to get a final Water Quality Index (WQI). The coefficients of variation of WQI over time did not exceed 0.43, indicating that water quality remained relatively stable over the considered period. The WQI theoretically ranges from 1 to 6, but here it ranged from 1 to 2.52 only (mean 1.35), indicating that all stations showed good to very good water quality.

**Distribution of main chemical components concentrations in the 47 river stations.**

Red vertical bars indicate concentration thresholds delimiting water quality classes, following the French implementation of the European Water Framework Directive 2000/60/EC (Ministère chargé de l'environnement (2013) « Guide technique actualisant les règles d'évaluation de l'état des eaux douces de surface de métropole », Annexe 4 : « Etat écologique des cours d'eau - Paramètres physico-chimiques généraux »). All river stations showed good to very good water quality when considering concentrations in nitrogen compounds (ammonium, nitrate and nitrite). Only one river station (MILEgl) showed mediocre water quality when considering concentrations in phosphorus compounds (total phosphorus and phosphate), suggesting local pollution. Nevertheless, it is noteworthy that 35% of monitored stations (n = 15) showed total phosphorus concentrations higher than 0.075 mg/L and may thus be considered as eutrophic[29].

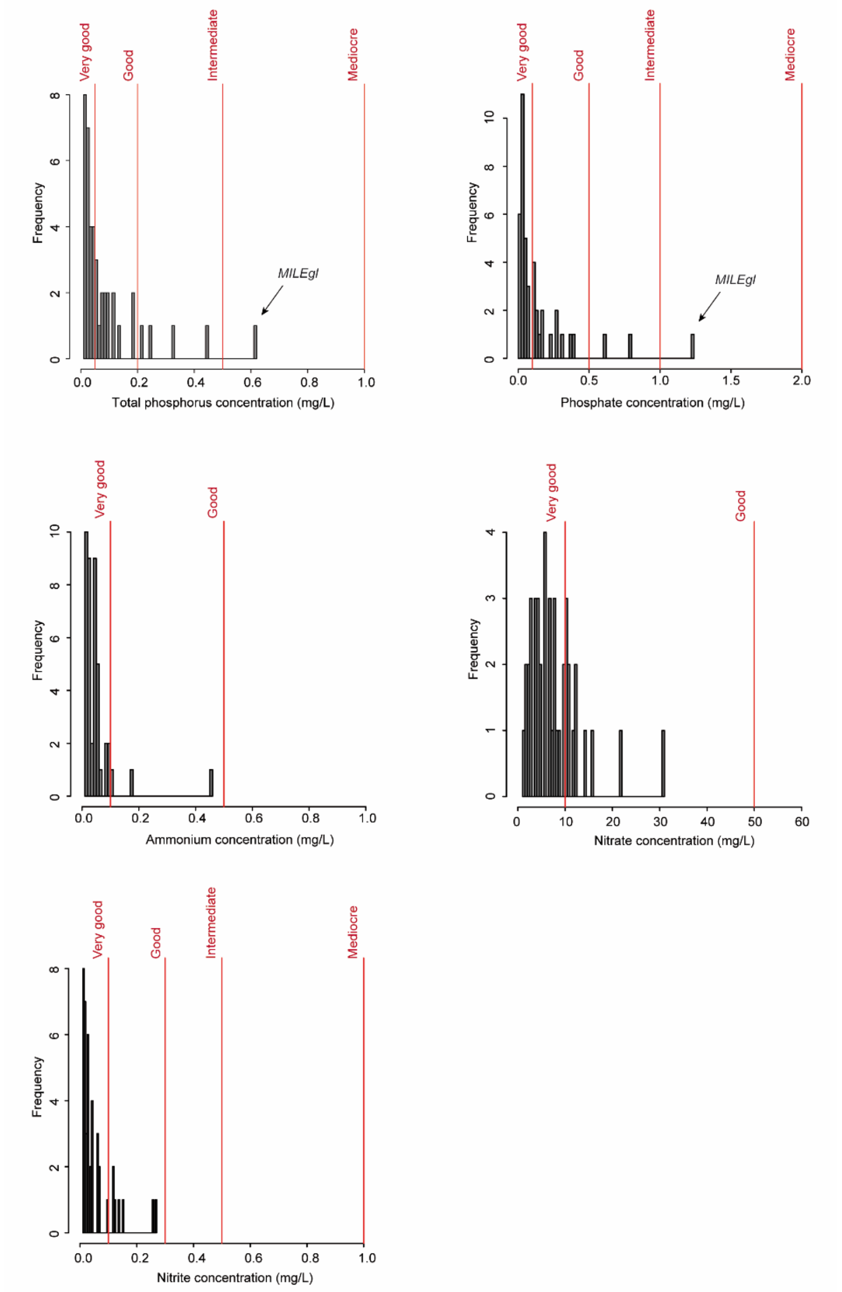

**Characteristics of the two first principal components (PC) based on environmental data.**

The following figure provides a visual representation of eigenvalues (corresponding to the amount of the variation explained by each PC; panel A) and of the two first PC accounting for 80.5 % of the total variance in environmental variables, with PC1 standing for the upstream-downstream gradient (fresh upstream stations on the one hand (negative coordinates) and warmer downstream stations on the other hand) and PC2 standing for the eutrophic gradient (nutrient-impoverished river stations (very good ecological status) on the one hand (negative coordinates) and nutrient-rich stations (medium ecological status) on the other hand; panel B). The percentage of variance explained by each component is also indicated. DFM: Distance from the river mouth; DFS: Distance from the river source; WQI: Water Quality Index.

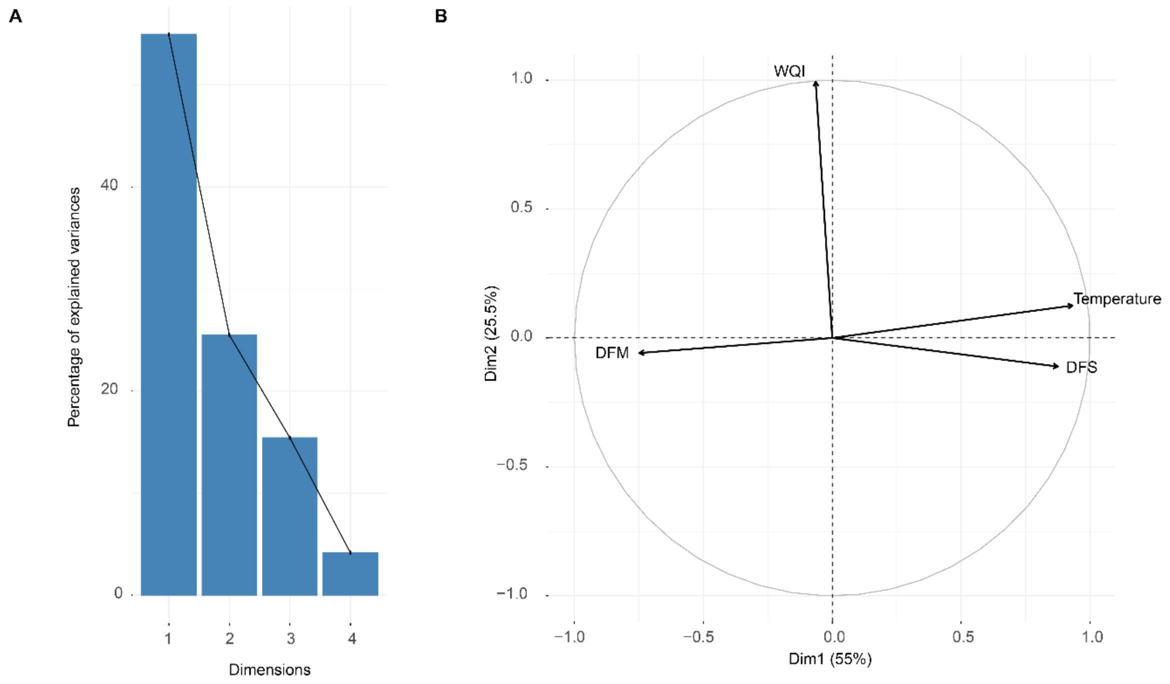

**Supplementary Note 6.** Initial causal models.

Initial causal graphs depicting all the investigated links among environmental (purple), bottleneck (blue), genetic (light green), and biomass variables (per capita and total; yellow). Mean biomasses are investigated in A, biomass stability in B. Links of interest (between IGD variables and biomass variables) are in bold.

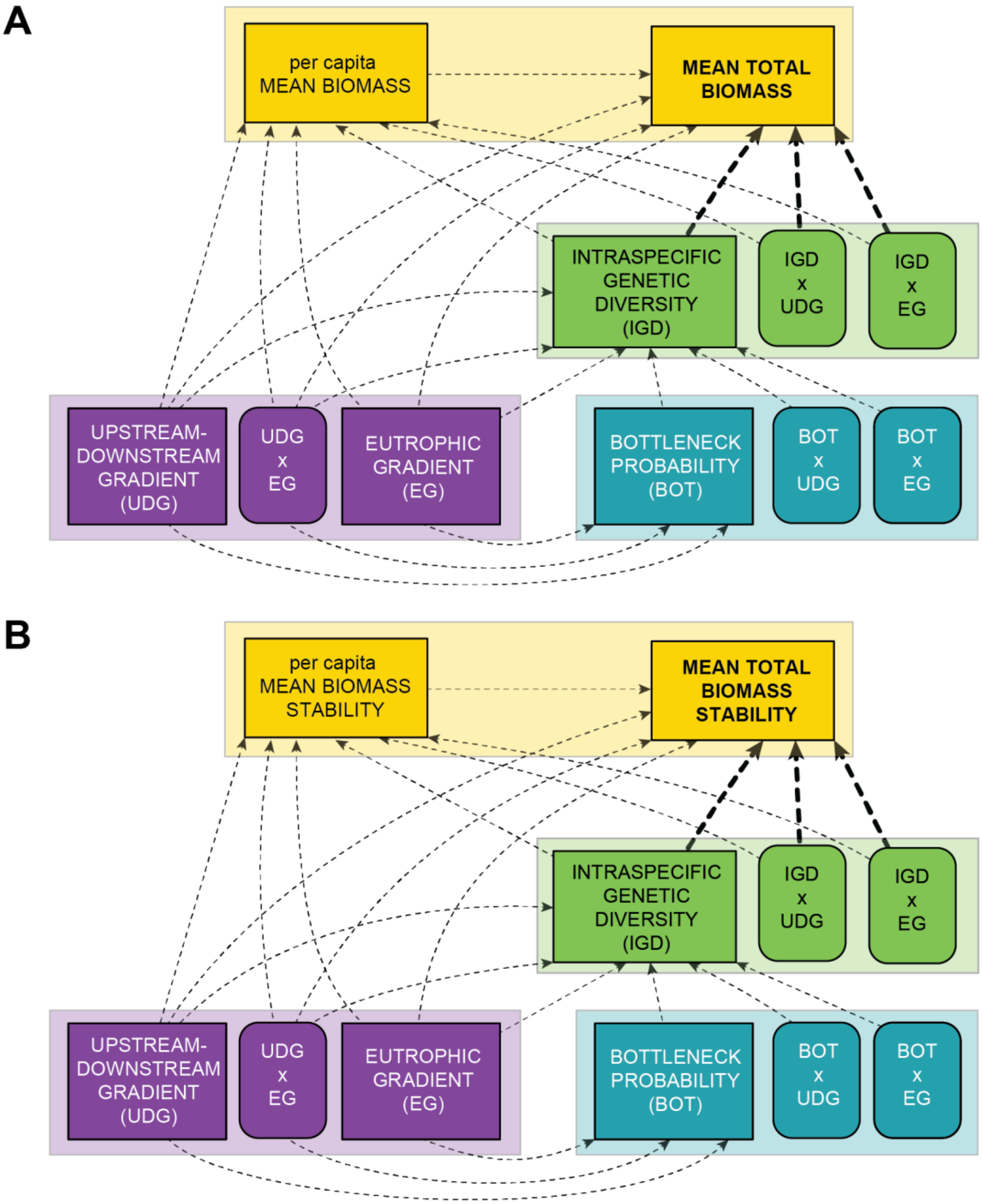

### REFERENCES


1. Poulet, N., Beaulaton, L. & Dembski, S. *J. Fish Biol.* **79**, 1436–1452 (2011).
2. Irz, P. *et al. Knowl. Manag. Aquat. Ecosyst.* 25 (2022) doi:10.1051/kmae/2022021.
3. Haegeman, B. *et al.* http://biorxiv.org/lookup/doi/10.1101/085852 (2016) doi:10.1101/085852.
4. Leigh, D. M., Hendry, A. P., Vázquez-Domínguez, E. & Friesen, V. L. *Evol. Appl.* **12**, 1505–1512 (2019).
5. Exposito-Alonso, M. *et al.* https://www.biorxiv.org/content/10.1101/2021.10.13.464000v2 (2021).
6. Aljanabi, S. M. & Martinez, I. *Nucleic Acids Res.* **25**, 4692–4693 (1997).
7. Futschik, A. & Schlotterer, C. *Genetics* **186**, 207–218 (2010).
8. De Kort, H. *et al. Mol. Ecol.* **27**, 2193–2203 (2018).
9. Catchen, J., Hohenlohe, P. A., Bassham, S., Amores, A. & Cresko, W. A. *Mol. Ecol.* **22**, 3124–3140 (2013).
10. Zerbino, D. R. & Birney, E. *Genome Res.* **18**, 821–9 (2008).
11. Li, H. & Durbin, R. *Bioinformatics* **25**, 1754–1760 (2009).
12. Li, H. *et al. Bioinformatics* **25**, 2078–2079 (2009).
13. Barnett, D. W., Garrison, E. K., Quinlan, A. R., Stromberg, M. P. & Marth, G. T. *Bioinformatics* **27**, 1691–1692 (2011).
14. Kofler, R., Pandey, R. V. & Schlötterer, C. *Bioinformatics* **27**, 3435–3436 (2011).
15. Rousset, F. *Mol. Ecol. Resour.* **8**, 103–106 (2008).
16. Rice, W. R. *Evolution* **43**, 223–225 (1989).
17. Van Oosterhout, C., Hutchinson, W. F., Wills, D. P. M. & Shipley, P. *Mol. Ecol. Notes* **4**, 535–538 (2004).
18. Fourtune, L., Paz-Vinas, I., Loot, G., Prunier, J. G. & Blanchet, S. *Freshw. Biol.* **61**, 1830–1845 (2016).
19. Jombart, T., Devillard, S. & Balloux, F. *BMC Genet.* **11**, 94 (2010).
20. Prunier, J. G., Dubut, V., Loot, G., Tudesque, L. & Blanchet, S. *Freshw. Biol.* **63**, 6–21 (2018).
21. Jombart, T. *Bioinformatics* **24**, 1403–1405 (2008).
22. R Development Core Team. (2014).
23. Belkhir, K., Borsa, P., Chikhi, L., Raufaste, N. & Bonhomme, F. (2004).
24. Garza, J. C. & Williamson, E. G. *Mol. Ecol.* **10**, 305–318 (2001).
25. Leblois, R. *et al. Mol. Biol. Evol.* **31**, 2805–2823 (2014).

26. Nikolic, N. & Chevalet, C. *Evol. Appl.* **7**, 663–681 (2014).

27. Paz-Vinas, I., Quéméré, E., Chikhi, L., Loot, G. & Blanchet, S. *Mol. Ecol.* **22**, 3279–3291 (2013).

28. Pella, H., Lejot, J., Lamouroux, N. & Snelder, T. *Géomorphologie Relief Process. Environ.* **3**, (2012).

29. Smith, V. H., Tilman, G. D. & Nekola, J. C. *Environ. Pollut.* **100**, 179–196 (1999).

30. Duffy, J. E., Lefcheck, J. S., Stuart-Smith, R. D., Navarrete, S. A. & Edgar, G. J. *Proc. Natl. Acad. Sci.* **113**, 6230–6235 (2016).

31. Oehri, J., Schmid, B., Schaepman-Strub, G. & Niklaus, P. A. *Proc. Natl. Acad. Sci.* **114**, 10160–10165 (2017).

32. Lefcheck, J. S. *Methods Ecol. Evol.* **7**, 573–579 (2015).

33. Fourtune, L. *et al. Am. Nat.* **191**, 491–508 (2018).

34. Aiken, L. S., West, S. G. & Reno, R. R. (SAGE Publications, 1991).

35. Paz-Vinas, I., Loot, G., Stevens, V. M. & Blanchet, S. *Mol. Ecol.* **24**, 4586–4604 (2015).

36. Grace, J. B. (Cambridge University Press, 2006).

37. Batista-Foguet, J. M., Coenders, G., Saris, W. E. & Bisbe, J. *Metodološki Zv.* **1**, 163–184 (2004).

38. Burnham, K. P. & Anderson, D. R. (Springer, 2002).

39. Hu, L. & Bentler, P. M. *Struct. Equ. Model. Multidiscip. J.* **6**, 1–55 (1999).

40. Rosseel, Y. *J. Stat. Softw.* **48**, (2012).
